# Exploration biases how forelimb reaches to a spatial target are learned

**DOI:** 10.1101/2023.05.08.539291

**Authors:** AC Mosberger, LJ Sibener, TX Chen, HFM Rodrigues, R Hormigo, JN Ingram, VR Athalye, T Tabachnik, DM Wolpert, JM Murray, RM Costa

## Abstract

The brain can learn to generate actions, such as reaching to a target, using different movement strategies. Understanding how different variables bias which strategies are learned to produce such a reach is important for our understanding of the neural bases of movement. Here we introduce a novel spatial forelimb target task in which perched head-fixed mice learn to reach to a circular target area from a set start position using a joystick. These reaches can be achieved by learning to move into a specific direction or to a specific endpoint location. We find that mice gradually learn to successfully reach the covert target. With time, they refine their initially exploratory complex joystick trajectories into controlled targeted reaches. The execution of these controlled reaches depends on the sensorimotor cortex. Using a probe test with shifting start positions, we show that individual mice learned to use strategies biased to either direction or endpoint-based movements. The degree of endpoint learning bias was correlated with the spatial directional variability with which the workspace was explored early in training. Furthermore, we demonstrate that reinforcement learning model agents exhibit a similar correlation between directional variability during training and learned strategy. These results provide evidence that individual exploratory behavior during training biases the control strategies that mice use to perform forelimb covert target reaches.

## INTRODUCTION

In arm movements like reaching to a glass of water, shifting gears while driving a car, or playing drums, the goal is to precisely reach a particular target in space. In such spatial reaching movements, different movement strategies may be used to get to the target location. One strategy is to move in a certain direction for a set distance, as in a feedforward movement (1). Another strategy is to move toward the target location based on the current location or sensory state of the limb (2, 3), as in a feedback movement (4, 5). The primary motor cortex is known to be crucial for targeted reaching movements across species (6-12) and it has been found to generate activity tightly related to reaching direction (13-18), as well as to integrate sensory feedback into new motor commands (19, 20). Despite the importance and universality of reaching actions, in which different strategies can achieve the same outcome (21), it is poorly understood what influences which strategy is learned and how they are controlled by the brain.

It has been proposed, that how a learner explores the task space during training determines what is learned about an action (21). It is thought that skilled actions are learned by assigning credit to the movement preceding the successful outcome (22, 23). Specifically, repeated credit assignment with practice allows the brain to gradually converge on what movement aspects were causal to success and reduce the variability of these aspects in a process called refinement (22, 24-27). Accordingly, when humans or non-human primates learn to reach to rewarded target locations, the pathlength or X-Y variance of reach trajectories are refined (28, 29). Thus, when different movement strategies can lead to the same outcome, which type of movements are explored during training could determine what can be assigned with credit and thus influence or bias which strategy is learned. How individual animals explore may depend on their previous experience (30-32), motivation, fatigue, or innate differences. Hence, confronted with the same task, individuals may learn to used different movement strategies based on what movements they explore during training.

Whereas past work has quantified refinement of relevant movement aspects during learning, this does not resolve which strategy was learned, as assigning credit to either strategy would likely reduce the variability of the movement in similar ways. The content of what was learned needs to be specifically probed. One way is to devise a test as has been widely used in the field of learning theory. Probe tests have allowed researchers to distinguish simple stimulus-response learning (33) from more cognitive place learning (34-36) by opening new paths or placing an animal at a new entry of a maze. And in instrumental learning, reward devaluation has been used to dissociate habitual (stimulus-response) learning from goal-directed learning (37). A series of studies in humans have devised similar tests to dissociate whether learned reach adaptations were represented in an intrinsic (joint space) or extrinsic (Cartesian space) reference frame by challenging participants with novel start positions or posture changes (38-42). Early studies in non-human primates showed that reaching a visible target from trained locations did not require proprioception, but from new locations it did (43, 44), suggesting that even reaches to visible targets can be performed using different control strategies.

In real-life scenarios, the spatial target of a movement is often not visible. In the example of the skilled drummer, the target of the arm reach doesn’t have to be visible and is merely a memorized location in space. It remains poorly understood how such reaches to hidden spatial targets are learned, and whether exploration influences what is learned. To causally test the role of specific circuits and neuronal cell types in these mechanisms, the mouse model provides untapped potential.

Here, we developed a novel forelimb Spatial Target Task (STT), where mice can freely explore the workspace and learn to move a joystick into covert targets, and where we can measure the refinement of forelimb trajectories, and dissociate whether a direction or endpoint strategy was learned. Mice are rewarded for entering a hidden circular target area in the workspace by moving a joystick from a set start position, similar to previous experiments in humans (28, 45). We refrained from using behavioral shaping or movement restriction, and positioned mice in a perched posture that enhanced exploration of the workspace. Mice discovered different targets in separate blocks and learned to successfully reach the targets. With training, we found significant refinement in both the initial direction of their movements and the targeting of the trajectory towards the target area. We also show that the spatial directional variability in joystick trajectories and the direction control is dependent on the sensorimotor cortex contralateral to the moving limb. By changing the start point in a small number of catch trials, we dissociated that some animals displayed direction learning and others endpoint learning, and show that the spatial exploration during learning biases which strategy animals learned to reach targets in space. Finally, we trained reinforcement-learning agents to perform the task and found that—as for the mice—exploration during training biased the agents toward either direction- or endpoint-based behavior.

## RESULTS

### Perched mice explore the workspace and learn to move a joystick into covert spatial target areas

We implemented the Spatial Target Task (STT) with a selective compliance articulated robot arm (SCARA) joystick as it provides a homogeneous horizontal workspace (46-48) and used a vertical manipulandum (49) that the mouse can hold and move around with one hand (Fig. 1a/b). Training started with a short pre-training period during which touching of the joystick was rewarded in the first phase, and forward movements of 4 mm in a 60 degree segment were rewarded in the second phase. At the end of pre-training, we defined target locations for each animal based on the mean direction of rewarded forward movements (Fig. 1d). The same target was rewarded for several days until high performance was reached (target training) (Fig. 1e). Mice self-initiated movements from a set start position and explored the workspace with complex trajectories (attempts) that were rewarded when they entered the hidden target area (hit) or ended either after a maximum time per attempt elapsed (7.5 sec) or if the animal let go of the joystick (200 msec, miss) (Fig. 1c). This structure allowed for self-paced and unrestricted exploration of the workspace when the joystick motors were disengaged, and no cues signaled where the target was.

**Figure 1.**
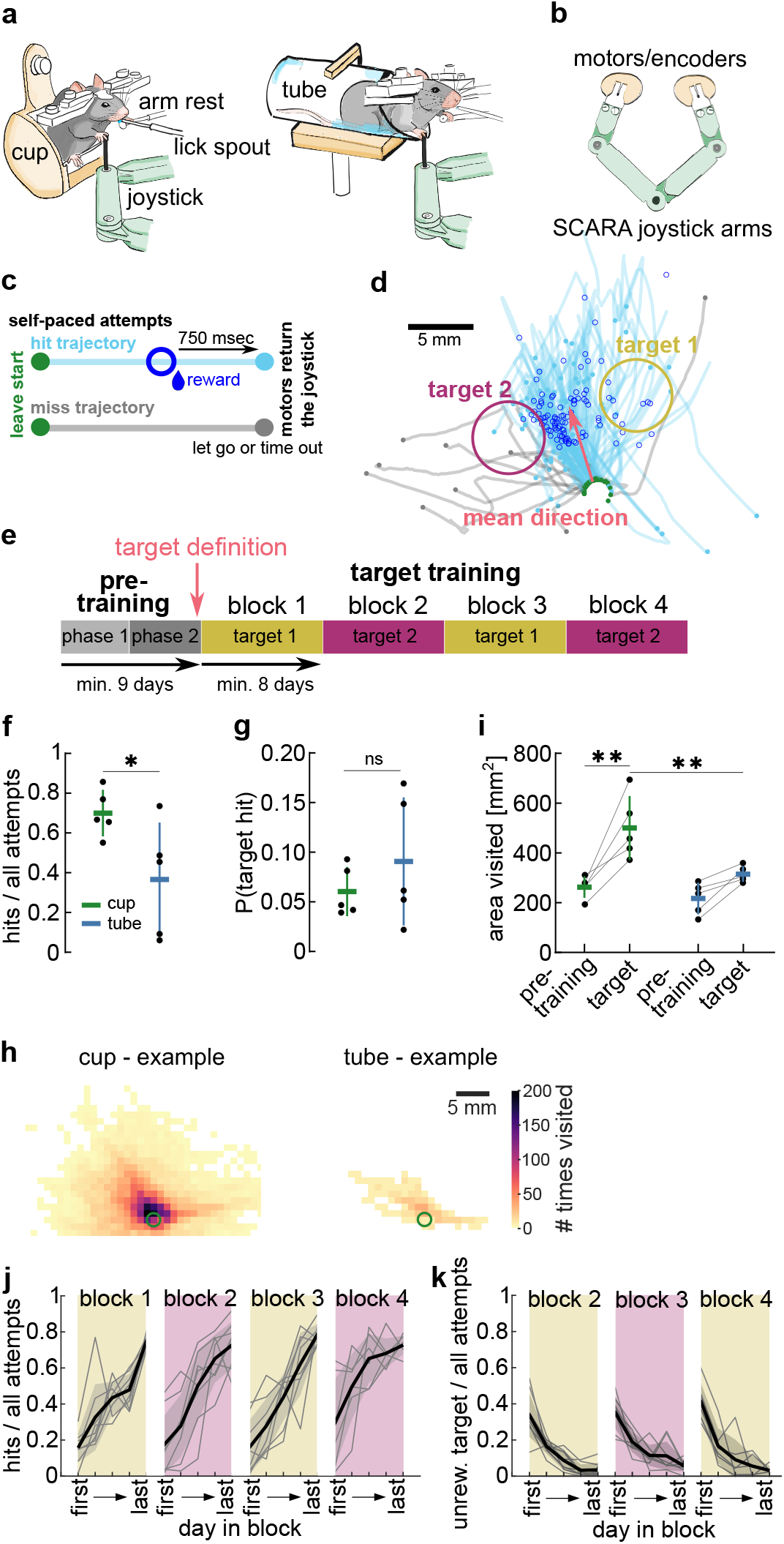
Perched mice explore the workspace and learn to move a joystick into covert spatial target areas. **a**. Schematic drawing of head-fixed mice using their right forelimb to move a SCARA (Selective Compliance Articulated Robot Arms) joystick while holding on to an arm rest with their left forelimb. Left: mouse sitting in the ‘cup’ assuming a perched posture. Right: mouse in an acrylic ‘tube’ assuming a quadrupedal posture. **b**. Schematic drawing of the SCARA joystick design showing a top view of both SCARA arms and motors/encoders to control and record the position of the joystick at the link joint. **c**. Schematic illustrating selfpaced attempts and resulting joystick trajectories. Every movement out of the start position (green dot) is an attempt. Hit trajectory: attempt enters the target area, a reward is dispensed (blue circle), and after a 750 msec delay, the motors move the joystick back to the start. Miss trajectory: target is not entered within 7.5 sec or the joystick is let go for >200 msec, motors move the joystick back to the start. **d**. Target definition. Example trajectories from the last pre-training session. Target locations defined 40° to the left and right of the mean hit direction (Suppl. Fig. 1a/b/h). **e**. Experimental design of Spatial Target Task (STT). Pre-training phase 1: touching of the joystick rewarded (4 days), phase 2: forward pushes rewarded. Target definition on the last day of pre-training. Target training: targets are rewarded in consecutive blocks of 8 days minimum per target. **f**. Ratio of hits achieved of all attempts made on the last session of block 1 of animals in the cup or tube. **g**. Probability of entering target 1 with all attempts made on the last day of pre-training. **h**. Example heat maps of visits to 1 mm2 bins in the workspace for an animal in the cup and the tube on the first day of the target block 1. The green circle indicates the start position. **i**. Total area visited by all attempts on the last day of pre-training and the first day target raining for animals in the cup and the tube. **j**. Ratio of target hits achieved on 5 equidistant days within each block from first to last day (selected days). **k**. Ratio of entering the previously rewarded target on selected days of blocks 2 - 4. f/g/i. n = 5 animals per group. j/k. n = 8 animals. All data shows mean +/-stdev and single animals. * p < 0.05, ** p < 0.01, ns: p > 0.05.

Exploration of the workspace is crucial to discover the target area and learn from reinforcement. In contrast to previous studies (11, 49-53) the reward contingency and target size were unchanged throughout the block such that only exploration would lead to target discovery, and no shaping was used. We also did not restrict the movement with the joystick, otherwise commonly done in mice (53-55), thus animals could move at any speed or stop mid-trajectory as often as they wanted. To further encourage animals to explore the workspace with their forelimbs, we tested whether head-fixing mice in a perched posture, as during food handling (56), would increase the forelimb range of motion compared to the commonly used quadrupedal positioning in a horizontal tube (11, 49, 57-60). We developed a custom cup-shaped holder (cup) that allowed animals to sit on their hindlimbs. Directly comparing animals trained on a target in either the cup (n = 5) or a standard tube (n = 5) (Fig. 1a), we found that the hit ratio reached was significantly higher in animals trained in the cup (Fig. 1f, unpaired t-test: t(8) = 2.42, p < 0.05, Suppl. Fig.1 c/d). This difference was not due to a difference in target locations between groups (Suppl. Fig. 1a/b, cup x/y: 0.32/7.48 +/-0.43/0.04 mm, tube x/y: 1.78/7.16 +/-1.55/0.60 mm, unpaired t-test, x: t(8) = 2.03, p > 0.05, y: t(8) = 1.19, p > 0.05). Animals of both groups showed low hit ratios at the beginning of target training (Suppl. Fig. 1a/b), and did not display a higher probability of entering the target after pre-training (Fig. 1g, unpaired t-test: t(8) = 0.98, p > 0.05)). In order to test whether animals in the cup were better able to explore the workspace with the joystick increasing their probability of discovering the target, we analyzed the area of the workspace visited by all trajectories of a given session. Heatmaps show an increased area visited on the first day of the target training for an example animal trained in the cup and an example animal trained in the tube on the same joystick rig (Fig. 1h). To quantify this exploration, we counted the total number of unique spatial bins visited by animals on the last day of pre-training and the first day of target training (Fig. 1i, Two-way ANOVA rep. meas., group: F(1,8) = 8.99, p < 0.05, day: F(1,8) = 32.57, p < 0.01, day x group effect: F(1,8) = 5.67, p < 0.05). There was no initial difference in exploration of the workspace between groups (Bonferroni corrected multiple comparison, pre-training day: t(16) = 0.94, p > 0.05), but cup animals significantly increased exploration to find the target on the first day of target training (cup group day comparison: t(8) = 5.72, p < 0.01), exploring a larger area of the workspace than animals in the tube (target day: t(16) = 3.83, p < 0.01). These findings suggest that mice in a perched posture were able to display a wider range of forelimb movements with the SCARA joystick (Suppl. Fig. 1e/f) and modulated their exploration to discover a hidden target area.

We therefore decided to use the cup for all other experiments and next tested whether animals could discover and learn different target locations, allowing for repeated measurement of learning and refining of skilled movements in the same animal. We pre-trained a cohort of 8 mice and defined two individual target locations for each animal (Fig. 1d, Suppl. Fig. 1h), which we rewarded in alternating blocks such that each target was repeated twice (Fig. 1e). Each block lasted until the performance criterion was reached and all animals progressed through each block within a maximum of 30 days (Suppl. Fig. 1i/k). During the first block, target 1 (lateral to the animal’s body axis) was rewarded in all animals. To combine data of animals with different block lengths we evaluated the performance on 5 equidistant days (from first to last) within each block. We discovered a steady increase in performance, with mice reaching high hit ratios (0.75 +/-0.07) at the end of each block, and a strong decrease after each target switch (0.19 +/-0.17) (Fig. 1j, Mixed-effects model, day in block: F(2.1,14.8) = 54.90, p < 0.01, block: F(2.0,13.8) = 3.08, p > 0.05). On the first day after a target switch, animals showed perseverative behavior continuing to enter the target of the previous block which decreased significantly as they explored the workspace and discovered the new target (Fig. 1k, Mixed-effects model, day in block: F(2.0,14.2) = 69.64, p < 0.01, block: F(1.5,10.2) = 1.81, p > 0.05).

These findings indicate that animals repeatedly learned to move the joystick into different target areas. However, the hit ratio only gives information about how frequently the target was entered, or how many times the correct action was achieved, but not how the exploration of the workspace or joystick trajectories evolved through learning (Fig. 2a).

**Figure 2.**
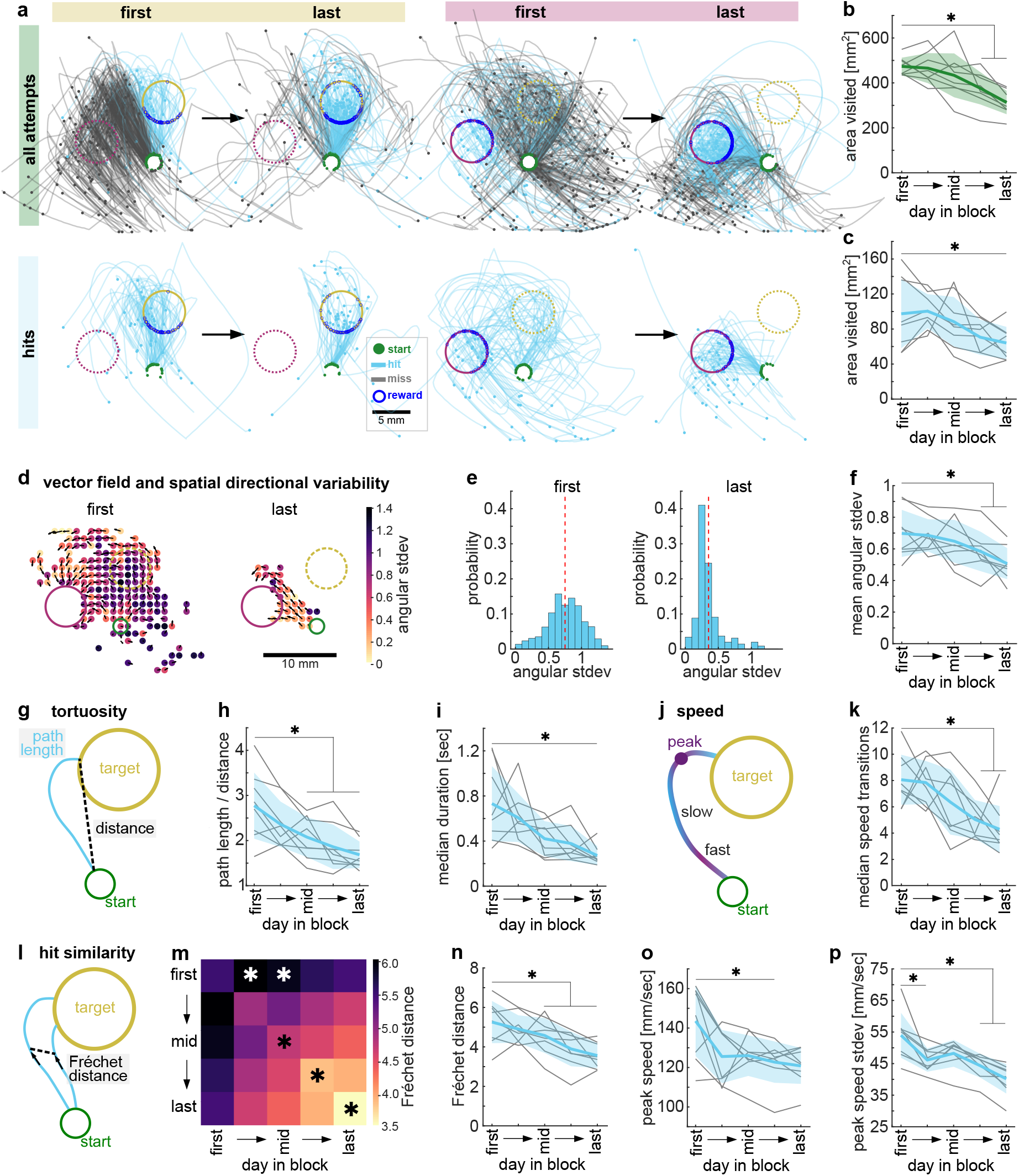
Mice explore the workspace with high spatial directional variability and tortuous trajectories. **a.** Example trajectories of first and last days of a target 1 and a target 2 block. Top: Trajectories of all attempts made in that session. Bottom: 50 subsampled hit trajectories from the same session. b-p. Analysis using all full length trajectories (green) or only hit trajectories from start to entering the target (blue) on 5 selected days and averaged across all blocks (n = 8 animals). One-way ANOVA with repeated measures, asterisks show Dunnett’s multiple comparisons between the first day and all other days of p < 0.05. Mean +/-stdev and single animals. **b**. Total area explored by all trajectories. **c**. Total area explored by 50 subsampled hit trajectories before entering the target. **d**. Example vector fields showing mean direction of all trajectories in a given spatial bin of the workspace (black arrows) and heat map of directional variability in that same position (spatial variability) on the first and last day of a target 2 block. **e**. Same data as in (d), histogram of the directional variability weighted by number of visits to each bin (angular stdev, dashed line = mean across all bins). **f**. Mean angular stdev across all bins for all animals. **g**. Schematic of path length and Euclidean distance from start to target entry used to calculate tortuosity (path length/distance). **h**. Tortuosity of hits for all animals. **i**. Median duration from leaving the start to entering the target of all hits. **j**. Schematic of speed of the movement along the trajectory indicating fast and slow sequences and the peak speed. **k**. Number of acceleration/deceleration transitions per trajectory for all animals. **l**. Schematic of pair-wise Fréchet distance (FD) calculation quantifying the similarity of trajectories. Similar trajectories have a small FD. **m**. Average similarity between hit trajectories within and across sessions measured by pair-wise FD. Hit trajectories within the later sessions are more similar to each other than hit trajectories within the first session (diagonal, black asterisks), and hit trajectories in early and middle sessions are dissimilar to those on the first session (top row, white asterisks). **n**. Same as diagonal data in (m) showing all animals. **o**. Mean peak speed of hits for all animals. p. Variability of the peak speed.

### Mice explore the workspace with high spatial directional variability and tortuous trajectories

We measured workspace exploration by counting the total number of spatial bins that were visited by all trajectories on 5 selected days across all blocks. We found that early in each block a large area of the workspace was explored, which reduced as more hits were performed (Fig. 2b, one-way ANOVA, F(2.6,18.2) = 13.62, p < 0.01). However, this would also be the case if animals merely stopped producing far reaching miss trajectories, while the hit trajectories stayed the same and were more frequently performed. To test this, we first measured the space explored by the hit trajectories alone and subsampled them to the same number of trajectories per session. We found that hit trajectories occupied a larger area of the workspace at the beginning of the block than at the end, even when we only considered the path from the start to entry of the target, when the reward was delivered (Fig. 2c, one-way ANOVA, F(2.5,17.5) = 5.95, p < 0.01). This data suggests that exploratory trajectories that were rewarded at the target entry (hits) were refined with training.

We next focused the analysis on these hit trajectories and investigated how they changed with learning. We used two measures of spatial exploration and refinement. First, the spatial directional variability, or how variable the average movement direction was at each position of the workspace. Second, the space explored within a trajectory, or how tortuous a trajectory was. In order to analyze the spatial directional variability, we generated vector fields of the binned workspace from hit trajectories of each session and quantified the angular standard deviation of the mean vector of all trajectories passing through a given spatial bin (Fig. 2d). This vector field analysis showed that the spatial directional variability across the workspace was high early in each block and then significantly decreased with learning (Fig. 2d-f, oneway ANOVA, F(2.4,16.6) = 7.91, p < 0.01). Furthermore, this decrease in spatial directional variability was specific to the rewarded segment of the hit trajectories, as the spatial directional variability of all full length attempts did not decrease with learning (Suppl. Fig. 3g, one-way ANOVA, F(2.7,18.6) = 0.97, p > 0.05).

In order to measure how exploratory a single trajectory was, we used a metric of tortuosity, defined by the integrated path length of the trajectory divided by the Euclidean distance between its start and endpoint at the target (Fig. 2g). Hit trajectories were significantly more tortuous in the beginning of the block than at the end (Fig. 2h, one-way ANOVA, F(2.7,18.6) = 15.06, p < 0.01), with trajectories becoming straighter with learning. Accordingly, the time required from leaving the start to entering the target decreased significantly across the blocks (Fig. 2i, 0.73 +/-0.36 sec to 0.27 +/-0.09 sec, one-way ANOVA, F(2.0, 13.8) = 5.36, p < 0.05). Furthermore, we found that early in the block when trajectories were more tortuous, they had more acceleration/deceleration transitions and this jerkiness also decreased with learning (Fig. 2j/k, one-way ANOVA, F(2.7,19.0) = 6.64, p < 0.01).

In order to investigate how the hit trajectories were refined across a whole block, we next asked how similar the shape and position between trajectories were within and between sessions of a block. We calculated the discrete Fréchet distance (61) (FD) between pairwise trajectories, which only compares points in the forward direction of travel but disregards differences in speed (Fig. 2l). Early in the block, the hit trajectories within a session were not similar to each other but became more similar with learning (Fig. 2m/n, oneway ANOVA, F(2.7,19.0) = 13.82, p < 0.01). Furthermore, hit trajectories on the first day of the block were dissimilar to hit trajectories on other days in the block, most strongly to hits in the middle of the block (Fig. 2m, one-way ANOVA, F(3.0,21.2) = 5.28, p < 0.01), further showing that the overall shape of the trajectories changed with learning across days. Overall, we found that animals initially explored the workspace with trajectories that moved in variable directions at different points across the workspace, were tortuous and jerky, and dissimilar to each other. These aspects of the movements were then refined, reducing variability and producing straighter trajectories as animals were rewarded for entering the target and increased their hit ratio (Suppl. Movie 1). Surprisingly, however, the decrease in tortuosity and jerkiness was not accompanied by an increase in movement speed, but rather the peak speed achieved per hit actually decreased with learning as well (Fig. 2o, one-way ANOVA, F(2.1,14.8) = 5.73, p < 0.05). Furthermore, this decrease in peak speed was also accompanied by a decrease in its variability (Fig. 2p, one-way ANOVA, F(2.*2,1*5.3) = 15.05, p < 0.01), suggesting an increase in control over the peak speed that would allow precise endpoint targeting.

### The precision of initial movement direction and targeting accuracy increase with learning

Animals may learn different strategies to move the joystick into the target area, namely moving in a certain direction for a certain distance, or moving towards a specific endpoint location from any position. We therefore measured refinement in these movement aspects independently.

First, in order to measure refinement of movement direction, we analyzed the direction of the initial segment of the hit trajectories and compared them to the optimal direction that led straight into the target (Fig. 3a, α: absolute difference between mean initial direction and target direction). We found that, as animals learned, their initial movement direction significantly approached the optimal target direction (Fig. 3b, one-way ANOVA, F(1.5,10.7) = 6.61, p = 0.02, last day α: 14.3 +/-3.9 deg). Importantly, they also significantly decreased the variability of their initial direction (Fig. 3c, oneway ANOVA, F(2.2,15.7) = 8.96, p < 0.01).

**Figure 3.**
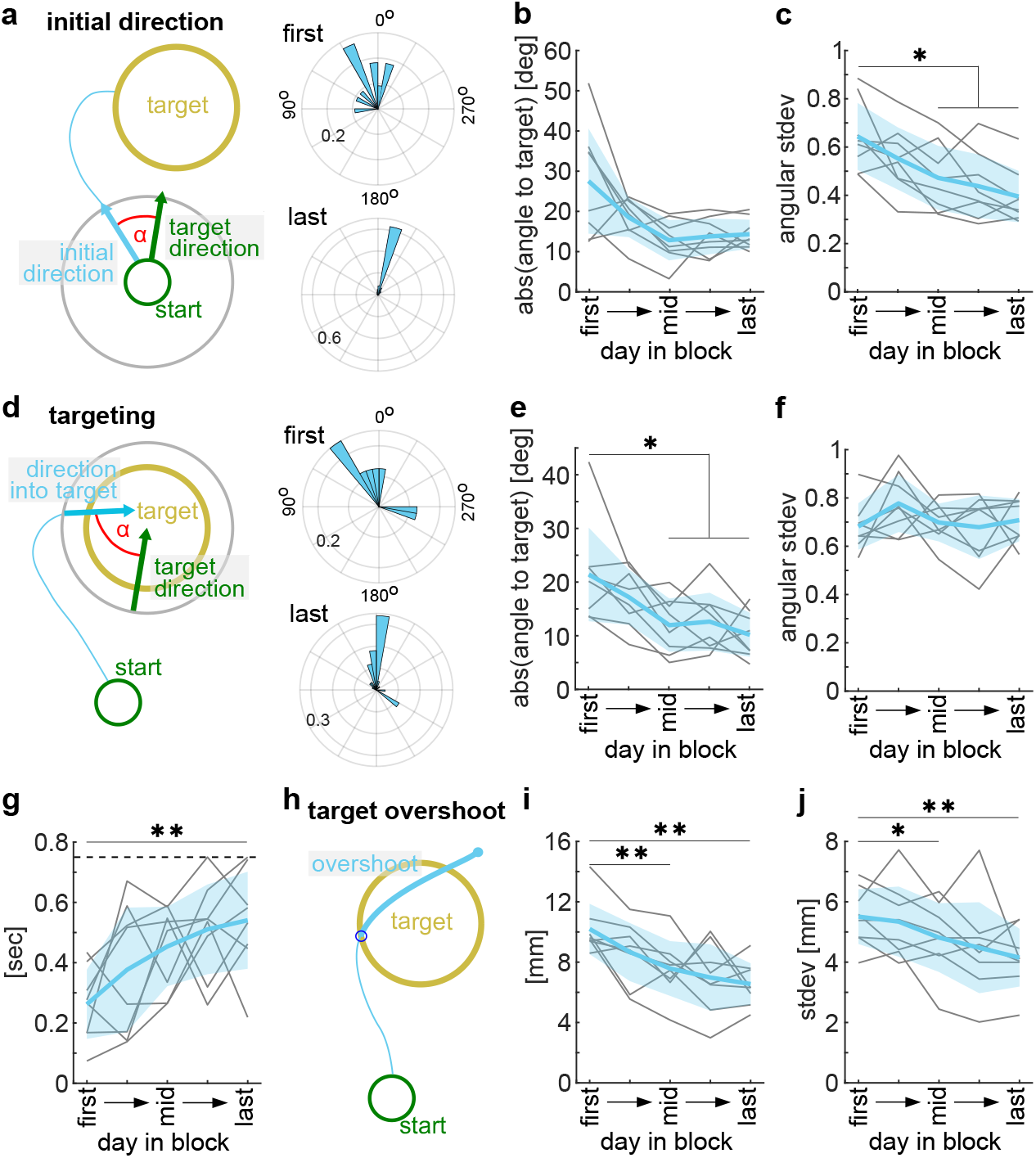
The precision of initial movement direction and targeting accuracy increase with learning. All analysis using hit trajectories on 5 selected days and averaged across all blocks (n = 8 animals). One-way ANOVA with repeated measures, asterisks show Dunnett’s multiple comparisons between the first day and all other days *: p < 0.05, **: p < 0.01. Mean +/-stdev and single animals. **a**. Left: schematic of initial trajectory direction calculation. Right: example polar histogram of initial hit direction on the first and last day of a block (probability). **b**. Mean initial direction difference to the direction straight into the target (0°). **c**. Variability of the initial direction. **d**. Left: schematic of direction of target entry calculation. Right: example polar histogram of the direction of target entry on the first and last day of a block (probability). **e**. Mean target entry direction difference to the direction straight into the target (0°). **f**. Variability of the target entry direction. **g**. Time spent in and closely around the target area after target entry until the delayed (750 msec, dashed line) returning of the joystick to the start. **h**. Schematic showing the overshooting of the target measured from the target entry point. **i**. The distance by which the target was overshot after entering. **j**. The target overshoot distance variability.

Next, in order to investigate the targeting behavior, we analyzed the final segment of the hit trajectory that led into the target circle. First, we measured the angle at which the target was entered taken from the point of passing through a concentric circle around the target with a 1 mm larger radius to the point of entering the target border and compared this direction to the straight direction from the start to the target center (Fig. 3d, α: absolute difference between mean direction into target and target direction). We found an overall approach to the target direction with learning (Fig. 3e, one-way ANOVA, F(2.8,19.4) = 7.36, p < 0.01, last day α: 10.2 +/-4.4 deg). Interestingly however, animals did not decrease the variability of target entry directions, even at high performance (Fig. 3f, one-way ANOVA, F(1.9,13.0) = 1.23, p > 0.05), suggesting a homing in on the target in the final segment of the reach and a correction of the movement if the target was missed with the initial movement direction. Furthermore, we found that animals dwelled significantly longer in and around the target area after target entry with learning (Fig. 3g, one-way ANOVA, F(3.2,22.1) = 5.04, p < 0.01) and the trajectory path that overshot the target entry point (Fig. 3h/i, one-way ANOVA, F(1.8,12.8) = 9.96, p < 0.01) as well as the over-shoot variability (Fig. 3j, one-way ANOVA, F(1.8,12.7) = 5.48, p < 0.05) decreased with learning.

These findings provide evidence that animals refined a fore-limb reach in a precise direction towards the target, but also shows features of endpoint-based control strategies with variable entry directions into the target. We next tested if these aspects of forelimb reaches were dependent on sensorimotor cortex in the mouse.

### A sensorimotor cortex stroke impairs movement direction and directional variability

Learning and performance of forelimb reaching movements has been shown to be dependent on sensorimotor cortex (7, 10, 62), but studies of rodents interacting with one-dimensional levers have found no impairment of learned skills upon motor cortex lesion (63). In order to test whether different aspects of spatial forelimb reaches are controlled by the mouse sensorimotor cortex, we performed confined cortex stroke lesions in expert animals. In animals trained to criterion in the STT (3-day average hit ratio > 0.65, n = 5), we performed a photothrombotic stroke (64) that caused a lesion of the sensorimotor cortex after animals had reached high performance in a target 1 block. We induced the lesion through a cranial window that we implanted over the left caudal forelimb primary motor cortex at the beginning of the experiment. We registered each brain volume and aligned it to the Allen Brain Atlas Reference Brain using BrainJ (65) to confirm the targeting of the lesions to the sensorimotor cortex for each animal (Suppl. Fig. 3c). The unilateral lesions encompassed a total volume of 9.9 +/-3.4 mm^3^ (Fig. 4a). The largest proportion of the lesioned volume was located in the primary (37 +/-5%) and the secondary motor cortices (27 +/-5%) (Fig. 4b). The lateral 17% and 10% of the volume also affected the primary somatosensory cortices of the forelimb and hindlimb, respectively (Fig. 4b). These strokes unilaterally lesioned 59 +/-7% of the total primary motor cortex, 40 +/-19% of secondary motor cortex as well as large parts of primary somatosensory cortices of the forelimb (82 +/-11%) and hindlimb (77 +/-7%) (Fig. 4c). Less than 12% of other somatosensory cortices and < 1% of striatum was affected by the lesions (Fig. 4b, Suppl. Fig. 3a/b).

**Figure 4.**
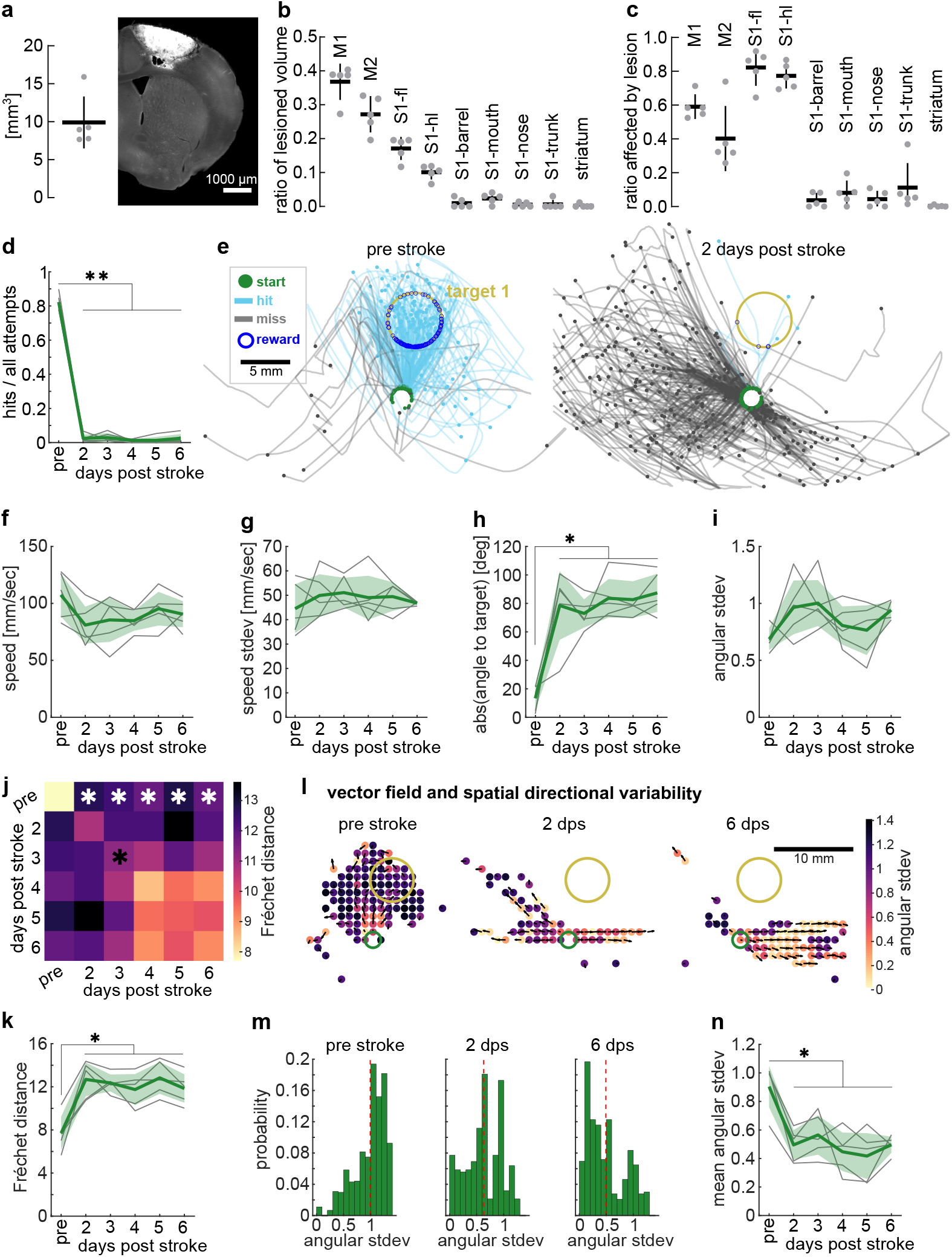
A sensorimotor cortex stroke impairs movement direction and directional variability. All analysis used trajectories from all attempts. One-way ANOVA with repeated measures, asterisks show Dunnett’s multiple comparisons between the pre-stroke day and all post-stroke days, *: p < 0.05, **: p < 0.01. Mean +/-stdev and single animals (n = 5). **a**. Left: total stroke lesion volume. Right: example histological coronal section showing lesioned brain region in auto-fluorescence. **b**. Relative volume of total stroke lesion that is part of different Allen Brain Atlas Reference Brain areas (sensorimotor cortex and striatum). **c**. Allen Brain Atlas Reference Brain areas that were affected by the stroke lesion showing the relative volume lesioned. **d**. Hit ratio before and acutely after the cortex stroke lesion. **e**. Trajectories from an example session before and 2 days after the stroke lesion (post stroke) of the same animal. **f**. Mean peak speed. **g**. Variability of the peak speed. **h**. Mean initial direction difference to target direction (0°) before and after the stroke lesion. **i**. Variability of the initial direction. **j**. Average similarity between trajectories within and across sessions measured by pair-wise Fréchet distance. Trajectories pre-stroke were more similar to each other than to trajectories post-stroke (top row, white asterisks), and trajectories poststroke were more dissimilar to each than trajectories pre-stroke (diagonal, black asterisk). **k**. Same as top row data in (j) showing single animals. **l**. Example vector fields showing mean direction of all trajectories in a given position in the workspace (black arrows) and heat map of the directional variability in that same position (spatial variability) of the session before the stroke and 2 and 6 days post stroke (dps). **m**. Same data as in (l), showing the spatial variability as histogram of angular stdev weighted by number of visits to a given bin (dashed line = mean angular stdev). **n**. Mean spatial variability across all bins before and after the stroke lesion. b/c. Abbreviations. M1: primary motor cortex, M2: secondary motor cortex, S1-fl: primary sensory cortex – forelimb, S1-hl: primary sensory cortex – hindlimb, S1-others: primary sensory cortex – other areas.

Animals were given a day to recover from the non-invasive photothrombotic stroke lesion and placed in the STT on days 2 through 6 post-stroke with the same target being rewarded as before the lesion. We found that the hit ratio after the stroke was strongly impaired with animals achieving almost no target hits (Fig. 4d, one-way ANOVA, F(1.6,6.4) = 614.7, p < 0.01). This was not due to a lack of task engagement as animals readily moved the joystick out of the start position to initiate attempts (Fig. 4e). To investigate how mice manipulated the joystick after the stroke lesion, we used markerless pose estimation on videos to track the joystick base and wrist of the mouse (Suppl. Fig. 3h/I, Suppl. Movie 2). We found that, as mice moved the joystick, their wrists were insignificantly closer to the joystick (Suppl. Fig. 3j/k, one-way ANOVA, F(1.9,7.5) = 3.38, p > 0.05) but the wrist placement in relation to the joystick was more variable (Suppl. Fig. 3j/l, one-way ANOVA, F(2.7,10.7) = 14.59, p < 0.01). Despite this deficit, the total number of attempts performed was not significantly different before and after the lesion with a small tendency for more attempts post-stroke (Suppl. Fig. 3d, one-way ANOVA, F(2.1,8.4) = 3.56, p > 0.05). However, as lesioned animals did not finish the session by reaching the maximum number of rewards, their sessions lasted longer and time between attempts (ITI) showed a tendency to be longer as well (Suppl. Fig. 3e, one-way ANOVA, F(1.5, 5,9) = 4.91, p > 0.05). Interestingly, the trajectories that animals produced after the cortex lesion were not slower, and they still reached a largely similar peak speed relative to before the stroke (Fig. 4f, one-way ANOVA, F(1.9,7.5) = 2.25, p > 0.05). Furthermore, the peak-speed variability was not affected (Fig. 4g, one-way ANOVA, F(2.2,8.7) = 0.51, p > 0.05).

Given the few hits achieved by post-stroke animals, we performed further trajectory analyses on all attempts made before and after the lesion to compare how animals moved the joystick. We first asked what effect the cortex lesion had on the movement direction and found that the mean initial trajectory direction was significantly farther away from the optimal target direction after the lesion (Fig. 4e/h, one-way ANOVA, F(2.4,9.7) = 17.71, p < 0.01) leading to a strongly impaired accuracy of trajectories. However, variability of the initial direction was not affected (Fig. 4i, one-way ANOVA, F(1.8,7.3) = 2.19, p > 0.05). To further investigate how consistent movements were before and after the lesion, we quantified the similarity between trajectories, again using the FD metric. To focus the analysis on the aimed movement into the target area, we only included the segment of the hits to the point of target entry and excluded any movements made during reward consumption.

We found that trajectories were very similar to each other before the lesion, but became dissimilar to each other early after the lesion (Fig. 4j, Suppl. Fig. 3f, one-way ANOVA, F(2.3,9.1) = 4.76, p < 0.05). Most strikingly though, the similarity analysis showed that trajectories produced after the lesion were very dissimilar to those before the lesion (Fig. 4j/k, one-way ANOVA, F(2.1,8.5) = 18.78, p < 0.01). Animals with a cortex lesion further seemed to perform center-out movements that extended to the end of the range of motion (Fig. 4e), without much spatial directional variability between trajectories. When we measured the spatial directional variability using vector field analysis during the STT of intact animals, we found that it was continuously high among full length trajectories of all attempts across days in a block (Suppl. Fig. 3g). After the stroke there was a strong and lasting impairment of this spatial directional variability (Fig. 4l-n, one-way ANOVA, F(2.1,8.3) = 13.75, p < 0.01).

Together, these results show that exploratory spatial variability and movement accuracy towards the target are dependent on sensorimotor cortex.

### A probe test reveals that individual animals learned to reach the target using different strategies

Expert mice had learned to launch their forelimb reaches more precisely in the direction of the target, but they still entered the target from variable directions at high performance. Thus from analyzing the refinement of the trajectories it remains unclear whether a direction or an endpoint strategy was learned. We next directly asked whether animals had learned a strategy of moving in a specific direction, or rather of guiding the movement to a specific endpoint location in space. We tested this with a probe experiment that challenged expert animals with new start positions.

After animals had reached the performance criterion on a target 1 block, we tested them in a session during which the joystick returned to novel start positions in a small number of catch trials. We defined two new start positions 40 degrees to the left or right rotated around the target with the same distance between start and target as the original start position (Fig. 5a). During each catch trial, the animal performed attempts from the new start for a maximum of 5 minutes or until the target was hit (to limit learning from reinforcement from the new start positions during the probe test).

**Figure 5.**
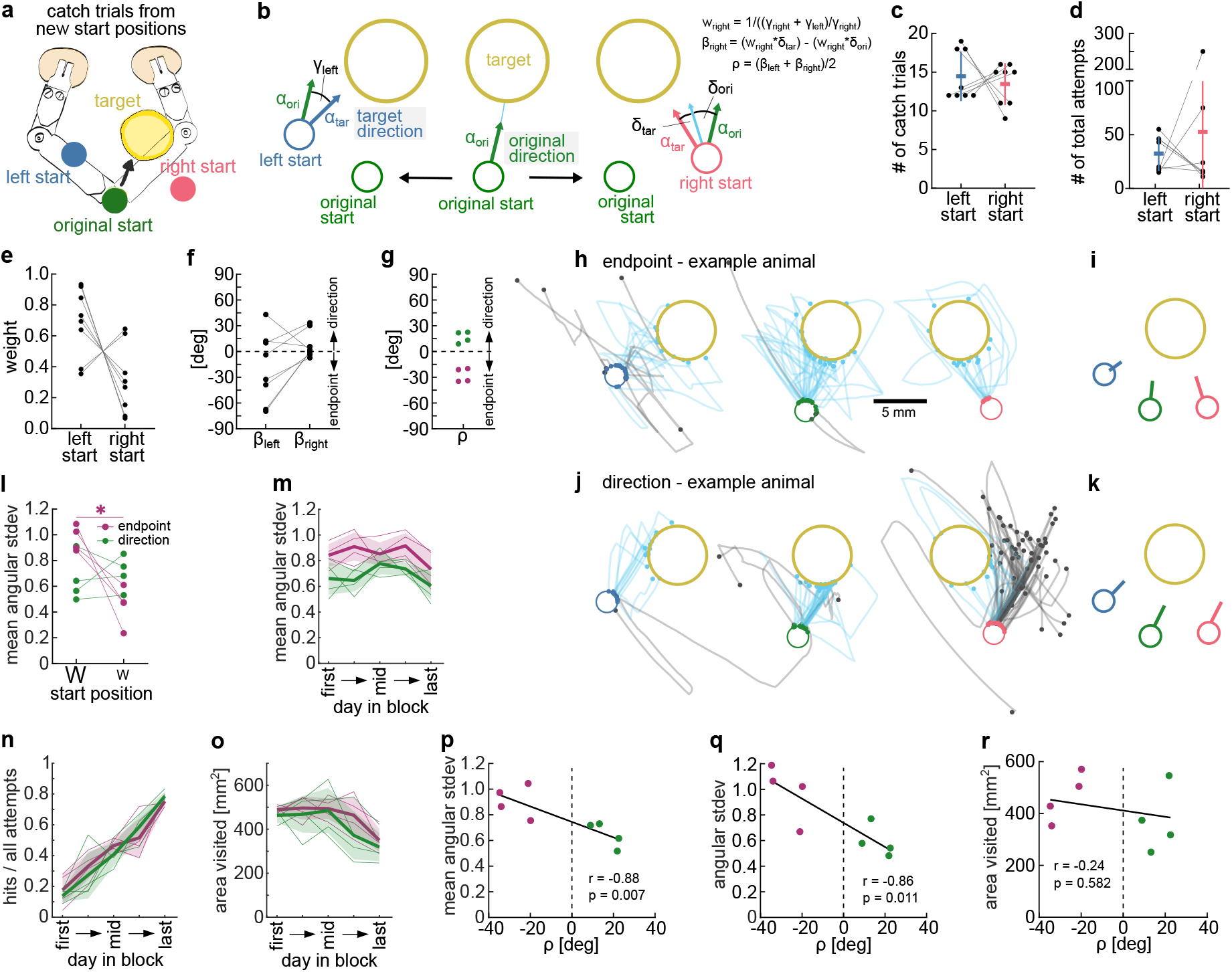
A probe test reveals that individual animals learned to reach the target using different strategies. All analysis used trajectories from all attempts made from the original or new start positions. Two-way ANOVA with repeated measures or t-tests were used. Asterisks show Dunnett’s or Bonferroni’s multiple comparisons, * p < 0.05, ** p < 0.01. Mean +/-stdev and single animals (n = 8 or n = 4). **a**. Schematic of start change probe test showing the new stat positions to the left and right of the original start position. **b**. Schematic of the initial vector analysis used to calculate the weighted angles (β) and final angle (ρ) to classify animals into ‘direction’ or ‘endpoint’ learner (see Materials and Methods for details). **c**. Total number of catch trials where the joystick moved to the left or right start for each animal. **d**. Total number of attempts performed from the new start positions during all catch trials per animal. **e**. Weighting factors for left and right starts determined from the difference angle (γ) between the original and target directions (α). **f**. Weighted angels (β) for both start positions. g. ρ angles for all animals. Animals with positive ρ, indicating mean initial directions closer to the original direction are shown green (direction learner). Animals with negative ρ, indicating mean initial directions closer to target direction are shown in maroon (endpoint learner). **h**. Example animal with negative ρ (endpoint learner) showing all trajectories from the new start positions and 40 subsampled trajectories from the original start position. Hit trajectories are only shown from the start to the target entry point. **i**. Average initial direction vectors for the same animal as in (h) showing vectors pointing to the target from new start positions. **j**. Same as (h) but for a direction learner animal. **k**. Same as in (i) but for the same direction learner animal showing initial vectors pointing in a similar direction from new start positions and original start. **l**. Spatial directional variability of attempts during catch trials from large and small weight start positions of endpoint and direction learners. **m**. Spatial directional variability of endpoint and direction learners during the target training (average of blocks 1 and 3, where target 1 was rewarded). **n**. Hit ratio of endpoint and direction learners during the same target training blocks. **o**. Workspace exploration measured in total number of bins visited of endpoint and direction learners during the same target training blocks. **p**. Same as (m) but showing variability of the early sessions in the blocks (selected day 2) and correlation to ρ angle used for classification. **q**. Variability of target entry angle of the early session in the block (selected day 2) and correlation to ρ angle used for classification. **r**. Same as (o) but showing area visited of the late session in the block (selected day 4) and correlation to ρ angle used for classification.

In order to dissociate whether an individual animal had learned a direction strategy or an endpoint strategy, we measured the average initial trajectory direction across all attempts made from each of the new start positions and compared it to the average direction from the original start position (original direction) (Fig. 5b). We classified animals that moved in a direction similar to the original direction from new start positions as direction learners and animals that moved in the direction of the target from new start positions as endpoint learners. We chose the initial vector direction as the basis for classification because it probes the immediate response to the new start location. The number of catch trials across all animals was 14.5 +/-3.2 for the left start and 13.5 +/-2.7 for the right start (Fig. 5c, paired t-test: t(7) = 0.48, p = 0.647). In all catch trials animals performed an average of 32.5 +/-16.5 attempts from the left start and 52.8 +/-82.4 attempts from the right start (Fig. 5d, paired t-test: t(7) = 0.62, p > 0.05). However, the median number of attempts from the right start was only 16.

The analysis of the initial direction vector also took into consideration how far the original direction was from the target direction of the new start position, i.e., how much the animal had to adjust their mean direction to hit the target from the new start, by weighting a larger adjustment more (weights, Fig. 5e). For all but 2 mice, the left start position was the more difficult start and thus had the larger weight. For both new start positions we then determined whether the initial direction of all attempts was closer to the original direction or to the target direction (Fig. 5f) and used the weighted average of that difference across both starts to classify the animal into endpoint or direction learner type (angle ρ, Fig. 5g). From the 8 animals that underwent the probe test, we found ρ > 0 for 4, which we classified as direction learners, and ρ < 0 for the other 4, which we classified as endpoint learners (Fig. 5g). Example trajectories from each learner type illustrate how the endpoint animal adjusted the trajectory direction early on in the attempt whereas the direction animal largely maintained the learned original direction (Fig. 5h/j). This overall difference becomes even more evident when looking at the mean initial-direction vectors of the same animals (Fig. 5i/k). We confirmed that there was no significant learning during the catch trials by comparing the hit ratio in later catch trials to the hit ratio in early catch trials of the session and found no change in hit ratio across the session in the endpoint (Suppl. Fig. 4a, mixed-effects model, catch trial: F(18,54) = 1.18, p > 0.05) or direction learner mice (Suppl. Fig. 4b, mixed-effects model, catch trial: F(17,51) = 0.83, p > 0.05).

Next we investigated how animals of the two learner types behaved during the catch trials in the probe test. Instead of separating their attempts by left or right start, we separated them into large and small weight starts, according to how much the animal had to adjust their original direction to hit the target from the new start, as before. We first tested whether animals were more successful in reaching the target from the new start positions than chance. For the start with a larger weight, we found no significant difference between learner types but a significant overall effect of chance showing that animals performed better than chance from the new starts (Suppl. Fig. 4c, two-way ANOVA rep. meas., chance: F(1,6) = 12.06, p < 0.05, learner: F(1,6) = 0.18, p > 0.05). For the smaller weight start we found the same chance effect but also a significant difference between learner types (Suppl. Fig. 4d, two-way ANOVA rep. meas., chance: F(1,6) = 10.11, p < 0.05, learner: F(1,6) = 8.52, p < 0.05). We also analyzed if animals adjusted their trajectories from the new starts enough such that they would yield a lower hit ratio if they were performed from the original start position. This analysis revealed that attempts performed from the new start positions were sufficiently different that they would have yielded significantly lower hit ratios from the original start (particularly in endpoint learners), with a significant interaction between start position and learner bias (Suppl. Fig. 4e, two-way ANOVA rep. meas., start position: F(2,12) = 22.49, p < 0.01, learner: F(1,6) = 12.68, p < 0.05, start x learner: F(2,12) = 6.95, p = 0.01). For the easier start, only endpoint learners adjusted the trajectories enough to lead to a significantly lower chance hit ratio.

We then focused our analysis on how animals were adjusting their movements from the new start positions. We measured the spatial directional variability by applying the vector field analysis to attempts from different start positions. Here we again found a significant interaction between start position and learner bias with endpoint learners showing a higher spatial directional variability from the difficult start than from the easier start, whereas direction learners displayed the same variability from both starts (Fig. 5l, two-way ANOVA rep. meas., start position: F(1,6) = 9.71, p < 0.05, learner: F(1,6) = 0.19, p > 0.05, start x learner: F(1,6) = 14.32, p < 0.01).

These results suggest that endpoint learners were adopting a strategy of increased spatial variability from the more difficult start position but not the easier one. This suggests that they had learned to move to the target position in space rather than a specific direction and could adjust their movement depending on the difficulty.

### The strategy learned is correlated with early spatial exploration

The endpoint and direction learning bias of different animals that we uncovered by the probe test could be the result of reinforcement of different movements during the target training. We next investigated whether animals that showed different strategies during the probe test had explored the workspace differently during training, biasing what they learned. We found that spatial directional variability across all attempts of blocks 1 and 3 (target 1 rewarded) was significantly higher in endpoint learners compared to direction learners (Fig. 5m, two-way ANOVA rep. meas., day in block: F(4,24) = 3.00, p < 0.05, learner: F(1,6) = 11.06, p < 0.05). This difference did not reflect a difference in performance as all animals reached high hit ratios with learning (Fig. 5n, two-way ANOVA rep. meas., day in block: F(4,24) = 45.27, p < 0.01, learner: F(1,6) = 0.11, p > 0.05). Furthermore, endpoint learners did not generally explore a larger proportion of the workspace, which would give them more experience with different positions in the space, as measured in the number of spatial bins visited (Fig. 5o, two-way ANOVA rep. meas., day in block: F(4,24) = 8.38, p < 0.01, learner: F(1,6) = 0.84, p > 0.05), but rather it was the way they explored the space that was more variable. The absolute difference of spatial directional variability between groups was largest early in the block (day 2 of the 5 selected days) and we used this day to test whether the degree to which each animal was classified a direction or endpoint learner (value of ρ) correlated with the spatial variability displayed during learning. The spatial variability correlated significantly with the learner bias as measured by the signed value of ρ (Fig. 5p, Spearman correlation, r = -0.88, p< 0.01). Additionally we found that animals that entered the target from more variable directions early in the block also showed a stronger bias of endpoint learning (Fig. 5q, Spearman correlation r = -0.86, p < 0.05). When we performed the same correlation analysis on the overall workspace exploration metric where the largest difference between groups during learning was on a late day in the block (day 4), we found no significant correlation between workspace visited and endpoint learning bias (Fig. 5r, Spearman correlation, r = -0.24, p > 0.05).

Together these findings suggest that animals that explored the workspace with more spatial directional variability between trials and entered the target from more variable directions were more likely to show an endpoint bias in the probe test. This could indicate that the reinforcement of variable trajectories may have led to the learning of an endpoint state because the common feature between rewarded trajectories was not a specific movement direction but an endpoint location. Conversely, for direction learners, trajectories were less variable, which could have allowed the reinforcement of a specific direction.

### Reinforcement-learning models confirm that exploration biases the type of learning

In order to probe whether this bias in what is learned could be the result of reinforcement of different exploratory behavior, we trained reinforcement learning model agents to solve a similar spatial target task, then confronted them with a start change probe test after learning.

In this model, a point-mass agent was trained to move through a continuous space from a start location to a hidden target location to obtain a reward (Fig. 6a). The agent was trained such that, given its state at each timestep, an action was chosen in order to maximize the sum of future rewards. The state consisted of a 5-dimensional vector, with two entries giving the agent’s Cartesian position, two entries giving the agent’s velocity, and one entry providing an “Go” signal, which is nonzero at the first timestep of each trial and zero otherwise. This Go signal was intended to model an internal signal within the brain to initiate movement, rather than an external sensory cue. Over the course of many trials, the agents learned to refine their trajectories, increasing their reward and decreasing the number of timesteps needed to reach the target (Fig. 6b). The trajectories of the agents, like those of the mice, were highly variable in early trials but much more refined in later trials (Fig. 6c).

**Figure 6.**
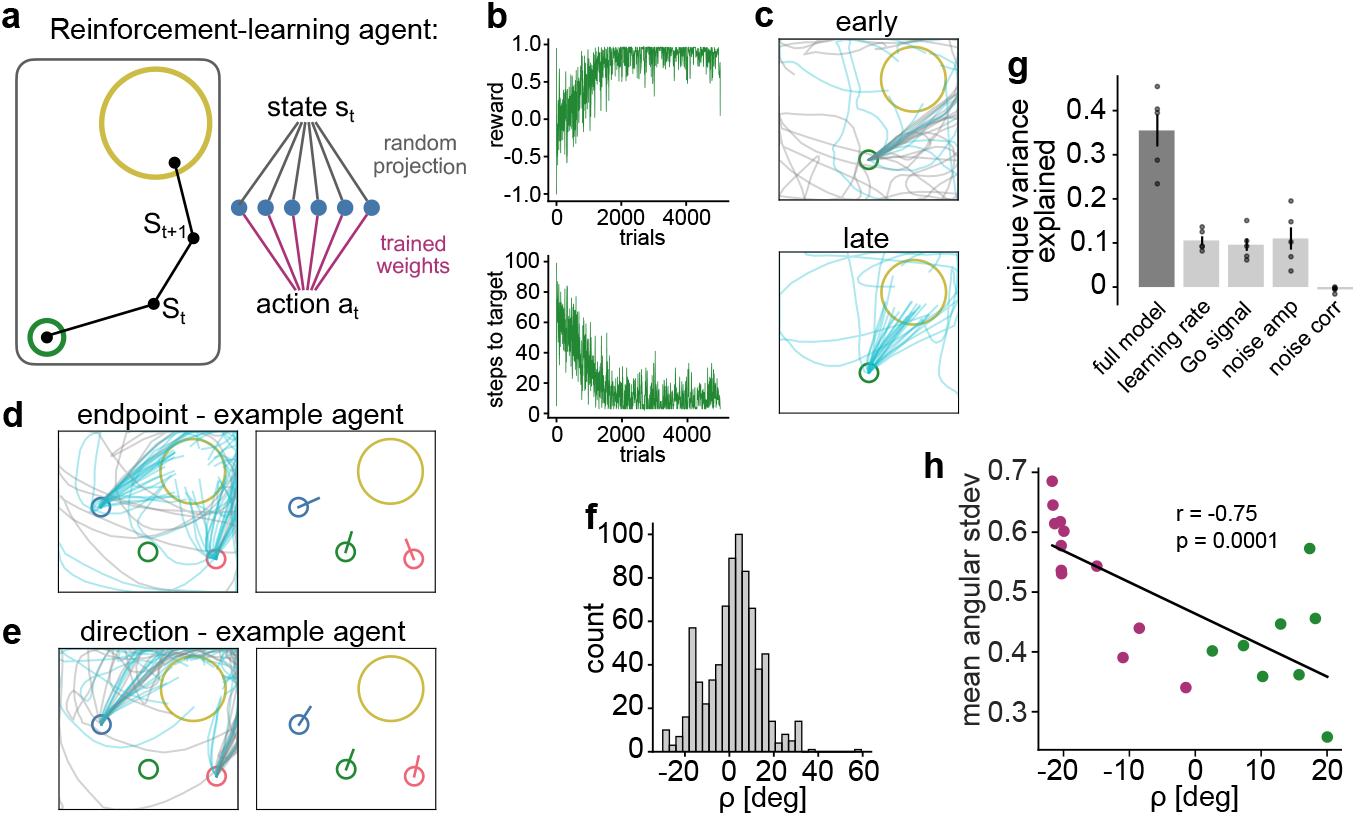
Reinforcement-learning models confirm that early exploration biases the type of learning. **a**. Single-layer network trained with policy-gradient learning to produce forces that move a point-mass agent to a goal location. **b**. Reward per trial (top) and number of steps required to reach the target (bottom) for an example agent throughout training. **c**. Trajectories for the same example agent, early (top) and late (bottom) in training. **d**. Trajectories and mean initial direction of an example agent that produced endpoint learner behavior. **e**. Same as (d) but for an agent that produced direction learner behavior. **f**. Distribution of ρ in an ensemble of agents trained with randomly chosen hyperparameters. **g**. The fraction of variance in ρ explained by a multivariate ridge regression model including all of the randomly chosen hyperparameters as regressors (full model), as well as the reduction in variance explained when leaving each of the hyperparameters out of the model (points correspond to different cross-validation splits of the data). **h**. The weighted spatial directional variability over training of n = 20 agents trained with identical hyperparameters but distinct initializations and correlation to their ρ angles.

In analyzing the experimental data, we found that, when the start position was changed, some mice exhibited direction learner bias, and others exhibited endpoint learner bias. Further, we had found that this difference between mice was correlated with the spatial directional variability of their trajectories during training. In order to test whether these same features and relationships would be exhibited by the model, we probed the trained agents by moving the start position at the end of training (Fig. 6d/e). We found that, as with the mice, the trained agents exhibited different performance types in response to the new start positions, with some exhibiting endpoint (Fig. 6d) and others direction learner (Fig. 6e) behavior.

Next, we asked which variables within the model most strongly affect the performance type of the trained agents. To investigate this, we randomly sampled the following hyperparameters for each agent: learning rate, amplitude of the Go signal, amplitude of the exploratory action noise, and correlation time of the exploratory action noise. This led to a mix of direction learner (ρ > 0) and endpoint learner (ρ < 0) agents (Fig. 6f). In order to examine which of these hyperparameters affect the performance type, we performed multivariate regression to predict ρ using the values of the four hyperparameters (Fig. 6g). The model captured a substantial fraction of the total variance in ρ, although less than half of the total variance was accounted for by these hyperparameters.

To quantify the unique contribution of each hyperparameter, we next performed the fit again using restricted models in which one of the four regressors was left out. We found that the learning rate, Go signal amplitude, and exploratory noise amplitude all accounted for roughly equal and mutually independent contributions, whereas the timescale of noise correlations accounted for essentially none of the variance in ρ (Fig. 6g). Interpreting our experiments in light of these results, we conclude that the first three of these quantities, if they differ across mice, may account for at least some of the differences in the observed learner types. However, most of the variance in ρ remained unaccounted for in these fits.

In our simulations, even agents with identical hyperparameters differed from one another in two respects: the randomly initialized weights (initial condition) and the random actions chosen at each timestep. We thus asked whether these two factors alone could be sufficient to account for the differences in performance type observed in our experiments. To test this, we trained agents with identical hyperparameters to perform the task and found that the observed range of ρ across agents was similar to that across agents with heterogeneous hyperparameters (Fig. 6h vs. Fig. 6f) and also to the distribution we found in mice (Fig. 5g). We next asked whether the performance type in these agents was related to the spatial directional variability of trajectories during training, as we had observed in the mice. We found that, as in the mice, higher levels of directional variability during training were associated with endpoint learner behavior, and lower levels of variability were associated with direction learner behavior (Fig. 6h, n = 20, Spearman correlation, r = -0.75, p < 0.01). Together, the results of our models provide further evidence that high levels of directional variability in the trajectories during training biases agents towards endpoint learning strategies.

## DISCUSSION

Here we introduced a novel spatial forelimb target task in which head-fixed mice learned to reach a circular target area from a set start position with a joystick in a 2D workspace. As we did not provide any ongoing feedback of the joystick position nor instructive visual cues of the target location, what was rewarded was the reaching of a covert location in space. This task can be solved by learning to move into a specific direction or to a specific endpoint location within the forelimb reference frame. We found that mice learned to accurately reach the target. As training progressed, they refined joystick trajectories by decreasing the spatial directional variability, tortuosity, speed, initial direction and approaching and overshooting the target. In order to test whether individual mice had learned a direction or an endpoint strategy, we implemented a probe test to dissociate what animals had learned. The probe test revealed that individual mice had a bias towards a direction or endpoint strategy that correlated with the spatial directional variability and target entry variability of their movements during training. Our results provide evidence that spatial exploration during learning influences the strategy used by individual animals to learn the task. In a reinforcement learning model, we found that agents trained to solve the STT showed endpoint and direction learning biases similar to mice. These biases were only partially explained by hyperparameters of the model and persisted when we fixed the hyperparameters. When we varied only the initial condition and exploratory action noise of different agents, the spatial directional variability during training was strongly correlated with the bias to show endpoint or direction learning, as in our mice.

These observations are consistent with studies suggesting that exploration during training is related to performance (21, 28, 66). Furthermore, our findings expand results from a human study where participants learned to move a virtual object to a target on a tablet and were then challenged with transfer tests during which there were small obstacles in the learned path (67). The study found that exploration during practice correlated with performance on the transfer tests and allowed participants to generalize to a new task-space. As in our mice, the correlation was not with the overall area visited but the trial-to-trial pattern of search (67). Here we showed that such exploratory patterns are correlated to the learning of distinct movement strategies in forelimb reaches.

Reinforcement of spatially variable trajectories, that also entered the target from different directions, would have biased the endpoint learner mice to assign credit to the target location in space and adopt a movement strategy that is more sensory feedback-based (reducing error to a desired endpoint) (43, 68). In direction learners, less variable trajectories, that entered the target from the same direction, would have been reinforced and led to learning of feedforward movement, potentially relying less on moment-to moment feedback (1, 5). As endpoint biased mice learned the task, sensory error-based learning (69-72) may thus have contributed to the refinement (73, 74) in addition to learning from reinforcement.

In our probe test, we classified animals into endpoint and direction learners based on their average initial directions from new start positions. However, we do not see these learners as binary, but rather as a gradient based on which strategy was reinforced more in individual animals. We thus restrict our conclusions to the average response, or bias, of an individual, but note that an animal may show direction or endpoint behavior in single attempts or even possibly switch strategies during an attempt in some cases (Fig. 5h/j). In humans, it has been shown that reaches to visible targets use direction-based control to move close to the target fast and then slow down to home in on it with endpoint-based control (68, 75).

We did consider that the variability of the joystick trajectories during training may have other causes than exploration (66, 76, 77). Variability can be due to noise in the sensorimotor system (78) that the motor system tries to reduce in the task-relevant domains (25, 79), and it could even impair learning (80). The remaining variability in joystick trajectories at high performance could be due to such noise. However, when animals had to discover a new target location at the beginning of the block, we saw an increase in variability of several movement aspects. Furthermore, during the probe test, in endpoint learner mice, the spatial directional variability was increased specifically in the more difficult new start location which required a larger adjustment of the direction. These findings suggest that variability was voluntarily increased for the purpose of exploration in all animals and to move into the **target from new start positions in endpoint learners (81, 82)**. Additionally, this indicates that endpoint learner animals may have improved error sensitivity as it has been reported that the movements made as corrections to sensory feedback errors can be a strong teaching signal that allows learning (83).

We also considered that the bias of showing direction and not endpoint responses during the probe test could have been confounded by sensitivity to the new position. It has been reported that mice reliably discriminate passive forelimb deviations of 2 mm (84), and our new start positions were ∼5.5 mm away from the original start. We are confident that animals learn this task using their proprioception for sensory feedback because animals performed the task in the dark and performance was unchanged when we removed their whiskers. Furthermore, there were no olfactory cues that could instruct where the target or start position was, as opposed to reaching tasks towards food pellets or water droplets (10, 85). In order to explore whether endpoint learner mice better integrated sensory feedback at new start positions, we measured whether they spent more time inside the ITI than direction learners, which would allow them to sense the new position for longer, but found no such relationship (data not shown). Yet, we did see attempts during which the movement direction was adjusted towards the target later in the attempt in direction learners, which could indicate that they used sensory feedback, but only once they were moving.

Our results showed that spatial target reaches are dependent on the sensorimotor cortex in mice. The motor cortex in rodents has been well established as a crucial structure for learning of forelimb movements in rodents (86-88). Particularly reach and grasp movements have been known to be dependent on sensorimotor cortex (7, 10, 12, 89). But the role of motor cortex in the performance of forelimb movements after their acquisition has been debated after a study in rats reported no effect of cortex lesions on a forelimb skill that required high temporal precision but less spatial precision than our task (63). Furthermore, deficits in reaching and grasping after sensorimotor cortex lesions have been considered to be mostly due to the reported loss of digit control. Accordingly, we found an increase in variability of the prehension of the joystick handle post stroke. Yet, animals were still able to make joystick movements and had specific deficits in their spatial directional variability and initial direction of reaches, whereas movement speed was not affected. These findings suggest a role for motor cortex in spatial target reaches (6, 9) but not in temporal control of movements (63, 90), as has been reported in primates (91). Interestingly, joystick movement directions after the stroke resembled the initial medial movements during pre-training, indicating a regression towards simpler movements without cortex. A similar phenomenon has been previously described after lesions to the thalamo-striatal projections in a task that required temporal precision (90).

In summary, our findings showed that in a spatial forelimb task in which a target location is reinforced, the spatial directional exploration of the task-space during learning affect what is assigned with credit and what is learned about the movement. This behavior platform in mice may be used to dissect the neural circuit mechanisms underlying exploration and the learning of directions or endpoints that lead to reward.

## METHODS

### Animals and experiments

All experiments and procedures were performed as approved by the Institutional Animal Care and Use Committee of Columbia University. Data of 21 male C57BL/6J wild type mice is shown. All animals were 3 -4.5 months old at the beginning of behavior training and maximum 8 months old at the conclusion of the experiment. All animals were single-housed after headbar implantation surgery and during the whole period of the behavior training.

10 animals were used to compare performance in the cup or tube (n = 5 per group). 8 animals were used in the main target training experiment with 6 animals being trained in 4 blocks, and 2 animals in 3 blocks. Cortex stroke lesions were performed in 5 of those animals. Data from an additional 3 animals is shown where we compare the performance before and after whisker trimming. These animals and the 8 animals used in the main target training experiment were also injected with retrograde adeno-associated virus (AAV) and implanted with cranial windows over the left forelimb motor cortex for use in a 2-photon imaging study. However, the main 8 animals were not used for 2-photon imaging because of insufficient viral expression or bone regrowth under the window and were instead used for behavior experiments according to 3R Animal Use Alternatives guidelines (USDA) to reduce the number of animals used in research. No difference in task performance was found between animals that were injected in different areas (Suppl. Fig. 1l/m).

### General surgical procedures and headbar implantation

Surgeries were performed under aseptic conditions. All tools were autoclaved and sterile surgical gloves were used during all procedures. Animals were anesthetized with Isoflurane (2% in oxygen). Buprenorphine SR was administered (1 mg/kg) before the surgery subcutaneously. The scalp was shaved using clippers and animals placed in a stereotaxic frame with cheek bars. Eye cream was applied and the skin cleaned with ethanol and Iodine ointment (Βetadine®) swabs. The scalp was removed using spring scissors and the skull cleaned and dried by applying 3% hydrogen peroxide and scraped with a scalpel blade. Dental cement (C&B Metabond Quick Adhesive) was applied to the exposed skull to build a cement cap and the metal headbar was implanted securely into the dental cement. Once the cement was dry the mouse was removed from the stereotaxic frame and placed into a clean cage on a heat mat and monitored until fully ambulating.

A custom headbar design was used with small U-shaped ends on either side of the straight cemented bar that allowed easy sliding in and securing of the mouse’s head in the head-fixation holders by tightening a screw through the U-shaped ends. The headbar implantation was combined with the cranial window surgery for animals that had windows implanted.

### AAV injections

Animals were injected with AAVretro(SL1)_Syn_GCamp6f (1.3e13 GC/ml, Janelia) or AAVretro_Syn_GCamp7f (1.0e13 GC/ml, lot #21720; 1.2e12GC/ml, lot #v52598, Addgene 104488-AAVrg) in the right dorsolateral striatum (DLS) (0.75 mm anterior, 2.55 mm lateral, 3.2 mm ventral of Bregma, 100 nl), or the right cervical spinal cord (segments C4-C7/8, 0.5 mm medial of central blood vessel, -0.9 to -0.5 ventral of surface, 60-75 nl per segment). Animal codes and injections are reported in Supplementary Table 1. All injections were made in a stereotaxic surgery through blunt glass pipettes mounted to a Nanoject (Nanoject III, Drummond). Injections in the DLS were made through a small craniotomy at the time of the cranial window implantation and the craniotomy sealed using superglue (Loctite Superglue Gel) and accelerant (Zip Kicker). Injections in the cervical spinal cord were performed in a separate surgery one week before the cranial window implantation. The animal was placed in a stereotaxic frame. A 1.5 cm skin incision was made from the back of the skull to the shoulder blades using a spring scissors. The acromiotrapezius muscle was cut sagittally along the midline for a few millimeters and the muscle retracted. The spinalis muscles were blunt dissected to gain access to the cervical vertebrae and the T2 thoracic process which was clamped to stabilize the spinal cord. The ligaments between the laminae were removed using forceps and the spinal segments injected through the intra-laminar space. The acromiotrapezius muscle was sutured using absorbable sutures (5/0) and the skin sutured with silk (5/0) sutures.

### Cranial window implantation

A 3 mm diameter biopsy puncher was used to mark a circle centered on 1.5 mm lateral/ 0.6 mm anterior of Bregma over the left hemisphere. The circle was carefully drilled and the bone removed using forceps. The craniotomy was cleaned with sterile saline and a glass window (2 circular coverslips 5 mm and 3 mm radius glued together using optical glue (Norland Optical Adhesive 63, Lot 201) was placed over the craniotomy and glued to the skull using superglue and accelerator. Dexamethasone (1 mg/kg) was administered after the surgery to prevent brain swelling.

### Sensorimotor cortex photothrombotic lesion

Rose bengal (Sigma Aldrich (330000)) was freshly diluted in sterile saline (10 mg/ml) and kept on ice and in the dark. Animals were anesthetized with Isoflurane and placed in a stereotaxic frame using cheek bars. The cranial window was cleaned with water and ethanol using q-tips and an opaque foil template with a circular hole of 3 mm diameter was placed over the cranial window. A cool light source (Schott KL 1600 LED) was attached to the arm of the stereotaxic frame and lowered on top of the cranial window. An intraperitoneal injection of 35 mg/kg Rose Bengal was performed and 6 min later the light source was turned on for 2-3 min at 3 Watt at 400 nm. After the light exposure the animal was removed from the stereotaxic frame and placed in a fresh cage. Rose bengal bioavailability was confirmed by inspecting the animals feces for a red tint on the next day.

### Stroke histology and reconstruction

Animals were anesthetized deeply with Isoflurane and transcardially perfused with 0.1M PBS and 4% Paraformaldehyde (PFA). Brains were extracted and post-fixed for 24 hrs in 4% PFA and then cut in coronal sections of 75 μm thickness on a vibratome. Sections were counterstained on slide using NeuroTrace 640 (ThermoFisher Scientific (N21483)) according to the manufacturer’s protocol, and then coverslipped in Mowiol. Brain sections were imaged on a Nikon AZ100 Multizoom slide scanner (Zuckerman Institute’s Cellular Imaging Platform) at 1 μm/pixel resolution using a 4x objective. Images from the slide scanner were aligned and registered to the Allen Brain Atlas Reference Brain using BrainJ (65). The aligned volumetric data was imported to Imaris 9 (Oxford Instruments) and the green autofluorescence that was acquired at 488/515 nm excitation/emission was used to manually delineate the lesioned area and render the total lesion volume. Custom Matlab (R2020a, Mathworks, Inc.) code was used to calculate the ratio of lesioned tissue for all affected areas of the Allen Brain Atlas Reference Brain.

### Video analysis of hand/joystick interactions after stroke

Videos of animals before and after the cortex stroke lesion were used to train a pose estimation model (92) to track 2 key points: the base of the joystick, and the wrist of the mouse’s right hand. The joystick key point was chosen at the bottom left corner of the joystick spacer which had good contrast for high-fidelity tracking. We labeled 1132 frames (1077 frames from prior cohorts and 55 frames from experiments reported here) across 51 sessions (44 sessions from 10 animals in prior cohorts and 7 sessions from 5 animals that received strokes in experiments reported here) using a custom labeler tool through amazon web service via Neuroscience Cloud Analysis As a Service (NeuroCAAS (93)) and trained a supervised model on Grid.ai.

Key point coordinates were analyzed by first removing all low-fidelity points with a likelihood of less than 0.99. Periods of active joystick movements in the x-dimension of the video frame were analyzed and the distance in x between the wrist and the joystick key points calculated for each animal and session. The pixel dimension was converted to mm for each video using a known distance in the frame close to the mouse’s hand (the front dimension of the cup). The calibrated average distance between the wrist and the joystick and its standard deviation was calculated for all animals and sessions. Since the wrist coordinate was subtracted from the joystick coordinate, positive distance values indicate that the wrist was located in front of the joystick (closer to the mouse’s body), and negative values indicate the wrist was located behind the joystick (further away from the mouse’s body).

### Joystick hardware and Spatial Target Task controls

The SCARA (Selective Compliance Articulated Robot Arm) joystick setup was built from commercial Thorlabs parts and custom-made acrylic or 3D printed pieces. Animals were head-fixed in either an acrylic tube (40 mm diameter, with a 45 degree angled opening at the front) or in a custom-designed 3D printed cup (copyright IR CU21353), and their headbar was positioned 23 mm above the tip of the joystick, offset slightly to the left and back for comfortable interaction of the right limb with the joystick. A metal screw was placed horizontally to the left of the joystick to be used as arm rest for the left limb. A lick spout (16 Gauge blunt needle) was placed in front of the mouth of the animal, connected through tubing to a solenoid (The Lee Company, LFVA1220310H) to dispense the reward. The solenoid also made an audible click sound upon opening acting as a reward signal. Animals were filmed from the right side at 30 Hz under infra-red light (USB Camera, 2.0 Megapixel, with a Xenocam 3.6mm, 1/2.7’’ lens). The SCARA joystick arms were 3D printed (Formlabs Form 2 resin) and manually assembled with shielded stainless steel ball bearings (McMaster-Carr, 7804K119) and shafts. The arms were linked at the front through a threaded shaft to which a metal M2 screw was mounted using a female spacer. The head of the screw served as the manipulandum for the mouse to hold and move the joystick with its hand. The 3D printed SCARA arms were attached to custom designed metal hinges that were mounted onto the shaft of the DC-motors which had integrated encoders (DC-MAX26S GB KL 24V, ENX16 EASY 1024IMP, MAXON Motors, Inc.). The motors were mounted on an acrylic platform which was positioned on a Thorlabs breadboard using Thorlabs parts. Dimensions of the SCARA arms were as follows: back arm length = 50 mm, front arm length = 35 mm, distance between motor shafts = 60 mm. A capacitive touch sensor was connected to the bottom of the joystick shaft and to the cannula of the lick spout to allow detection of joystick touch and licking.

The Spatial Target Task (STT) was controlled through a microcontroller board (Teensy 3.6, Arduino) and breakout board platform developed in-house by the Instrumentation Core at the Zuckerman Institute (TeenScience, github.com/Columbia-University-ZMBBI-AIC/Teenscience). All task designs were written using the Arduino IDE. The main task loop (2 kHz) recorded the encoder positions, calculated the Cartesian coordinates of the joystick through forward kinematics, and triggered task states depending on the joystick position and touch sensor inputs. Touch sensors were read at 40 Hz. The joystick was actively moved to the start position and maintained at the start position by a proportional integral derivative (PID) algorithm running via interrupt functions and also operating at 2 kHz. Angular positions, for the PID controller and calculation of joystick position, were measured from the encoders with a resolution of 0.09 degrees. This gave a resolution of 0.06 ± 0.02 mm for joystick position across the workspace (Suppl. Fig. 2a). To move the joystick to the start position using the motors, the angular position of both encoders was compared with the start position (in angle space). Differential values were calculated to produce a driving pulse width modulation signal which was sent to the motors via the H-Bridge power amplifiers included on the TeenScience board. This control signal was calculated and recorded in torque space. To implement an inter-trial-interval (ITI, see below), a maximum force threshold was set in torque space resulting in joystick endpoint force thresholds between 7 and 11g (Suppl. Fig. 2b/i/j). The mice were required to remain below the force threshold for 100 msec before starting a new attempt. The motors were disengaged and no forces were generated during the active exploration of the workspace by the mice. The joystick position and all task events were recorded via serial output commands through Bonsai (OpenEphys). With all of the closed-loop control being performed on the Teensy microcontroller, 4 setups were run on a single standard computer (ASUS, Intel i7 CPU, 16 GB RAM).

### Spatial Target Task design

Starting on the day before behavior training began, animals were food restricted overnight and then given an individualized amount of chow food after each training session to maintain their body weight at 80% of pre-training baseline. Each session lasted until a maximum number of rewards were achieved. But a maximum time of 150 min was allowed and sessions were ended earlier if the animal stopped making attempts or didn’t consume the reward for an extended period of time, which happened mostly during pre-training. The average duration of a pre-training session was 21 min, and a target training session 33 min. The reward delivered from the lick spout was a 10 ul drop of 7.5% sucrose in water.

#### Pre-training

Pre-training consisted of 4 days of Phase 1, and 5 or more days of Phase 2 (Fig. 1e). For both pre-training and the target training, animals could perform movements with the joystick in a self-paced uncued manner. The joystick was initialized at the beginning of the session at a fixed start position (0/65 mm from the motor axis, 1 mm radius). Once head-fixed, the animal could move the joystick out of the start position and explore the workspace without any force generated by the motors for 7.5 sec per attempt. In Phase 1 of pre-training, a reward was given independent of the joystick position upon initial touching of the joystick (with a delay of 500 msec on days 1 and 2 and 1000 msec on days 3 and 4) and at random intervals between 5 to 15 sec for continuous touching of the joystick. The session ended after 100 rewards were delivered.

For both Phase 2 and the target training, animals had to move the joystick out of the start position and explore the workspace to receive a reward. If the exploration of the 2D space did not lead to a reward within 7.5 sec or if the mouse let go of the joystick for > 200 msec, the attempt ended and the motors engaged and moved the joystick back to start (Fig. 1c, miss). If the criteria for a reward was met, a reward was delivered and after a 750 msec delay the motors engaged and moved the joystick back to the start (Fig. 1c, hit). In Phase 2 a reward was given for moving the joystick in a forward direction initially within a 100 degree then a 60 degree segment for at least 4 mm. The reward was delayed randomly between 15 to 50 msec upon reaching the required radius. The session initially ended after 50 rewards and as performance improved after 100 rewards. The animal progressed to target training if it reached a rewarded attempt ratio of > 0.5 at the 60 deg segment. Using the rewarded trajectories on the last day of Phase 2, 2 target locations were defined for each animal as follows (Fig. 1d). The mean initial direction of all rewarded trajectories was calculated and a target center defined 40 degrees to the left and right of the mean direction at 8 mm distance from the start position. The target radius was set at 2.75 mm. Supplementary Fig. 1a/b/h shows the target positions for all animals. The initial hit probability was not different between the two targets (Suppl. Fig. 1j, paired t-test: t(7) = 1.76, p > 0.05).

#### Target training

During any given block in the target training, one of the 2 defined targets was rewarded. If a joystick movement entered the target circle a reward was instantaneously delivered. We did not delay the reward as we found it impaired learning in our mice in pilot experiments. In a similar task in humans, delaying the reward even by a few 100 msec also severely impaired learning (94). Once the joystick was returned to the start at the end of a rewarded (hit) or unrewarded (miss) attempt, an inter-trial-intervaI (ITI) started during which the joystick was continuously held at the start position by the motors until the animal exerted less than 7-11 g (average 8.2 g) force against the joystick in any direction for a minimum of 100 msec (hold period) (Suppl. Fig. 2b/e/i/j). Animals were not required to touch the joystick during the hold period, but analysis of the touch sensor showed that they mostly did (Suppl. Fig. 2d/g/h). At the end of the hold period the motors disengaged and the animal was allowed to initiate a new attempt by moving the joystick out of the start position (post-hold period) (Suppl. Fig. 2c/d/f). No cue was given that the hold period had ended.

The first 2 sessions in each block were completed after 50 target hits were achieved, the next 2 after 100 hits, and the rest after 150 hits. Each movement of the joystick out of the start position was counted as an attempt and task performance was calculated as the number of attempts entering the target circle (hits) divided by all attempts (hit ratio). When an animal reached a 3-day average hit ratio of > 0.65, and had received at least 900 rewards over a minimum of 8 sessions, the target was changed in the next session and a new block began (performance criterion).

For the comparison between animals positioned in the cup or the tube, we also defined a termination criteria. If an animal was trained for 10 days on a target without ever exceeding a 3-day average of 0.1 hit ratio, or if an animal did not reach the performance criteria within 21 days, the block was ended. Three animals from the tube group reached the termination criteria and were consecutively switched to be trained in the cup as well for a within animal comparison (Suppl. Fig. 1e/f).

#### Start change probe test

The probe test was performed after performance criteria was reached on target 1 either in block 3 or in an additional shorter block 5. For each animal, two new start positions were defined by rotating the original start position (0/0 mm) 40 degrees to the left (left start) and right (right start) around the target center, such that the distance between start and target was maintained but the direction to the target was changed. In the probe test session, the original start position was used for the first 10 hits to allow the animal to settle into the session. Then the joystick returned to either the left or right new start position to begin the first catch trial. During the catch trial, the animal could make multiple attempts to hit the target but to prevent learning from reinforcement, the catch trial ended after the target was hit from the new start position and the joystick returned to the original start position for 5-10 hits before the next catch trial. The order of left and right start positions chosen for the catch trials was randomized. If no hit was achieved in a catch trial within 5 min, the catch trial was ended and the joystick returned to the original start position. The probe test was completed when the animal achieved 200 rewards.

### Additional behavior tests

Three additional animals that were trained in the STT under a 2-photon microscope were also used to test the requirement of whiskers for task performance. Animals were trained on a baseline day at expert level on target 1, after the training session animals were briefly anesthetized using Isoflurane and all their whiskers on both sides were cut to a length of about 2-3 mm using scissors. Animals were placed in their home cage to recover from the anesthesia and tested in the task again the next day.

### Analysis

All data was analyzed using custom Matlab code (matlab engine for python R2019b, Mathworks, Inc.) run from a Python analysis pipeline (Python 3.7.8) through a DataJoint database (datajoint 0.13.0) (95). For the main target training experiment, data of 5 selected days per block is shown. For each block that includes the first and the last day, and 3 equidistant days in between to span the whole block. Data was analyzed for all blocks and a mixed-effects repeated measures model was used to determine if there was a significant effect of the day in the block (p < 0.05), but not of the block itself (Suppl. Table 2). Only if there was no significant effect of the block itself (p > 0.05), the data was then combined by averaging the selected days across all blocks for each animal. Blocks with target 1 took more days to reach the criterion than blocks of target 2 (Suppl. Fig. 1k, Mixed-effects model, target: F(1,7) = 8.62, p < 0.05; Bonferroni’s multiple comparison: block 1 vs 2: t(7) = 2.96, p < 0.05, block 3 vs 4: t(5) = 3.35, p < 0.05), even when we disregarded the required minimal number of rewards or days (data not shown, Mixed-effects model, target: F(1,7) = 10.06, p < 0.05; Bonferroni’s multiple comparison: block 1 vs 2: t(7) = 2.97, p < 0.05, block 3 vs 4: t(5) = 3.82, p < 0.05). However we found no significant savings effect of target repetition in either cases (Mixed-effects model, block: F(1,7) = 4.29/4.32, p > 0.05).

One-way ANOVA with repeated measures and Dunnett’s correction for multiple comparison, as well as paired and unpaired t-tests were used for statistical analysis. A p-value of less than 0.05 was considered statistically significant, and p-values are reported as > 0.05 (not significant, n.s.), < 0.05, and < 0.01.

#### ITI force analysis

End-point forces of the joystick were calculated from the value of two analog signals recorded from the TeenScience board. These analog voltages were linearly related to the absolute torque generated at each motor. Calibration of the analog Volts to motor torques was performed using a single axis version of the joystick (with a single 35 mm link). The TeenScience board was programmed to simulate a simple un-damped spring, and the analog Volts associated with various weights were recorded (2, 5, 10, 13 g). The conversion factor from Volts to motor torque was obtained from a linear regression. Joystick forces for each data sample were then calculated from the calibrated motor torques using the forward dynamics of the SCARA.

The ITI was divided into 3 periods: pre-hold, hold, and post-hold. During the pre-hold and hold periods, the joystick was maintained inside the start circle by the motors. The pre-hold period continued until the force was below the 7-11 g threshold. Only pre-hold periods of 10 msec duration or more were analyzed and the average force was calculated across this window (Suppl. Fig. 2b/i). If the force on the joystick was below the threshold when the post-hold period began, the hold period was entered immediately. The hold period was defined as a 100 msec window during which the force on the joystick remained below the threshold. The average force during the hold period was also calculated and was below 1 g on average (Suppl. Fig. 2b/j). The post-hold period followed the 100 msec hold period, during which the motors were disengaged. The post-hold period ended when the animal initiated a new attempt, moving the joystick outside the start position. The probability of touch during the ITI was calculated by determining for each time sample whether the touch sensor was active or not and averaging across the entire ITI (3 hold periods; Suppl. Fig. 2d/g/h).

#### Preprocessing of trajectories

Trajectories sampled at 500 μsec intervals were downsampled to 6 msec intervals for ease of handling the data.

#### Trajectories used for quantification

For each attempt a joystick trajectory was recorded. To quantify aspects of refinement of the rewarded attempt, only trajectories that entered the target area (hits) were analyzed. Furthermore, of those hit trajectories, only the path from start to the point of target entry was used in the analysis, as movements performed during reward consummation and voluntary returning to the start position were not considered part of the reinforced movement. For all metrics of variability, 50 trajectories were subsampled for all sessions unless there were less than 50 trajectories available. Analyses on pre- and post-cortex stroke lesion sessions were performed on full length trajectories of all attempts, to assess the overall movement differences, and because the small number of hits post-stroke did not allow analysis of hits only. For trajectory similarity analysis pre- and post-stroke, hit trajectories from start to the point of target entry were used.

### Calculation of trajectory metrics

Area visited: For the quantification of the area explored, the workspace (40 × 40 mm centered around the start) was divided into 1 × 1 mm bins and for each trajectory the bins visited were calculated using the Matlab ‘histcount2’ function. Bin counts for all trajectories were summed up and binarized to discount dwell time per bin and multiple visits to each bin. The total number of visited bins is reported as the area visited. In the experiment comparing animals trained in the cup or tube, the analysis was limited to the workspace in the forward direction of the start position 40 × 23 mm as the workspace behind the start position was smaller for animals trained in the tube.

#### Tortuosity

For each trajectory the total path length was calculated and divided by Euclidean distance between the first and the last point of the path (Fig. 2e). For each session the average tortuosity was calculated.

#### Vector field analysis

The workspace was divided into 1 × 1 mm bins. For each trajectory the vector going from one bin to the next along the trajectory path was recorded and assigned to the corresponding bin. If the same trajectory passed through a bin more than once, a separate vector was recorded for each pass through. If the trajectory stopped inside the bin and then continued, the combined vector was recorded. For each bin, the vectors for all trajectories of a session were concatenated and bins with less than 3 vectors were excluded. For the remaining bins, the vector angles were calculated using the ‘atan2’ Matlab function. The mean vector angle and angular standard deviation (bounded between the interval [0, √ 2]) was calculated using the CircStat circular statistics toolbox (96) functions ‘circ_mean’ and ‘circ_std’. Mean angles and angular standard deviations were used to plot vector field and heat map figures. The bin-wise angular standard deviation was then weighted by the number of visits to that bin and the mean of these weighted values was calculated as an overall metric for spatial directional variability within a session.

#### Trajectory similarity

The Fréchet distance (FD) was calculated as a scalar measure of similarity between trajectories that considers only forward movement along the trajectory but does not require uniform speeds (Fig. 2l). For each animal, the trajectories for all selected sessions were concatenated and the discrete FD between all pairwise trajectories was calculated using the ‘DiscreteFrechetDist’ Matlab function (61). The FD is 0 for two trajectories that follow the same exact path even if their speed profile is different. For each animal and session the mean within session FD was calculated and compared across blocks. The FD of trajectories from different sessions within each block was also calculated and averaged across animals to show as heatmaps.

#### Speed metrics

For all speed metrics, the speed of the downsampled (6 msec interval) trajectory was smoothed using a 30 msec moving average (‘smooth’ function in Matlab). For analyzed hits, only the trajectory from the start until entering the target was considered. The maximum value of the smoothed speed was calculated for each trajectory and averaged across all trajectories of a session (peak speed) and the standard deviation calculated as well (peak speed stdev). The jerkiness of trajectories was calculated by counting the transitions between acceleration and deceleration (sign changes of d(speed)/d(t)) for each trajectory. For each animal and session the median number of speed transitions was calculated.

#### Duration

The time from leaving start to entering the target was taken for each hit trajectory and the median duration calculated per session.

#### Time spent at target

Time spent in and 1 mm around the target after the target was entered was calculated for each hit and the median per session calculated.

#### Target overshoot

The target overshoot was calculated as the pathlength between the point of the trajectory entering the target and the end of the trajectory, when motors engaged (750 msec after entry). For each session the standard deviation of the target overshoot was calculated as a metric of targeting variability.

#### Initial vector direction

The initial trajectory vector was defined from the point at which the trajectory left the start circle (1 mm radius) to the point of it crossing a circle of 3.75 mm radius from the start position center (Fig. 3a). The components of all vectors were averaged to calculate the mean vector per session. For each vector and the mean vector the angle was calculated using the ‘atan2’ Matlab function, subtracted from the angle leading straight to the target center, and the absolute value reported (initial direction). The angular standard deviation across all vectors was calculated using the ‘circ_std’ function (see ‘Vector field analysis’).

#### Vector of target entry

The direction at which the target was entered was calculated by taking the vector from the point of the trajectory crossing a circle 1 mm bigger than the target itself to it crossing the target border (Fig. 3d). The angle of this vector was subtracted from the angle leading straight from the start position to the target center and the absolute reported. The angular standard deviation was calculated as for the initial vector direction.

### Classification of direction and endpoint strategy

To assess the degree to which animals moved towards the target or in their learned original direction from the new start positions, attempts from the different start positions were analyzed separately to calculate initial vector directions. First, the initial direction of all attempts from the original start position was calculated as described earlier using a radius of 3.2 mm around the start center (original direction, α ori). For the left and right new start positions the angle of the vector pointing to the center of the target was also determined (target direction, α tar left and α tar right). Depending on the nature of the animal’s original direction, the range between the original and target directions was smaller or larger for the left or the right start position (Fig. 5b), so their absolute difference (γ = abs(α tar - α ori)) was calculated to determine a weighting factor w (see below). For all attempts an animal made from the new start position during all catch trials, the mean initial vector direction was determined. This direction was then subtracted from α ori and from α tar resulting in 2 absolute angle differences for each start position (δ ori and δ tar). These angles measure if the animal’s attempts from the new start position were performed more in the original direction (direction learner) or in the target direction (endpoint learner). The angles δ ori and δ tar were then multiplied by the weighting factor (w) such that the attempts from the start where original was further away from target were weighted stronger. w-left = 1 / ((γ left + γ right) / γ left). For each start, the weighted δs were then subtracted from each other (w*δ tar - w*δ ori) resulting in a final signed angle β for each start. To classify each animal into overall ‘direction’ or ‘endpoint’ learner, the signed βs from both left and right start positions were averaged resulting in a final angle ρ used in the correlation analyses. If ρ > 0, the attempts from the new start locations were overall closer to the original direction than the straight target direction, which was considered a ‘direction’ performance. If ρ < 0, the attempts from the new start locations were overall closer to the straight target direction than the original direction, which was considered an ‘endpoint’ performance.

### Reinforcement learning model

The model consisted of an agent trained with reinforcement learning to map a 5-dimensional state representation onto actions that maximize reward. To create a more useful state representation for the agent, the first 4 components of the state vector (the agent’s position and velocity) were randomly projected via untrained weights onto a set of 99 radial basis functions, and the last component of the state vector, which had amplitude A_Go_ in the first timestep and was zero in subsequent timesteps, was concatenated onto this state representation. The radial basis functions had Gaussian kernels of width (in the spatial dimensions) 1/4 times the width of the arena and (in the velocity dimensions) 1/16 times the width of this arena (where the timestep size relating position to velocity was Δt = 1). The 2-dimensional action then consisted of a linear readout from this basis via trained weights, added together with exploratory noise 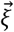. The noise was correlated from one timestep to the next and was given by the equation

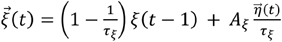

where *A*_ξ_ is the noise amplitude, is the noise correlation time, and *η*_*i*_(*t*) ∼ *N*(0,1).

The action consisted of a force applied to the agent, which influenced the agent’s velocity according to Newtonian dynamics. Specifically, the agent’s position at each timestep was updated as 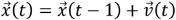, and the agent’s velocity was updated as 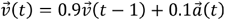. The arena boundaries were impenetrable, such that the agent could move along the boundary but not beyond it. The reward was -1/T for each timestep that the target area was not reached, and 1 if the target area was reached. Each trial was completed when the target area was reached or, if the target area was not reached, after T = 100 timesteps. To prevent the trivial solution in which the agent produces a very large action to reach the target in a single timestep, the agent received a negative reward contribution 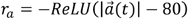.

The agent’s weights were trained using actor-critic learning with learning rate α to maximize the expected sum of future rewards, with temporal discount factor γ = 0.99. The critic learned the value associated with each state via a separate linear readout from the radial basis function representation. To facilitate credit assignment over multiple timesteps, eligibility traces were used in both the actor and critic, with time constants τ_e_ = 10.

In cases where the hyperparameters were chosen randomly, they were sampled uniformly from the range log_10_αϵ(−4,-2),

*A*_Go_ ϵ(0,10), log_10_*A*_ξ_ ϵ (−1,0.5), 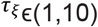. In cases where the hyperparameters were fixed, they were set to α = 0.001, *A*_go_ = 10, *A*_ξ_ = 2, and = 3. Only agents that met a performance criterion of reaching the target in 90% of the last 25% of trials were used for subsequent analysis.

In the multivariate regression model, the four hyperparameters listed immediately above were used as regressors in a leave-one-out ridge regression model using RidgeCV from the Python scikit-learn library. The model was fit on data from 80% of the trained agents and tested on the remaining 20%, and the results shown in Fig. 6 illustrate the test performance across the k = 5 possible data splits.

## Acknowledgements

We would like to thank Luke Hammond and Darcy Peterka from the Zuckerman Institute’s Cellular Imaging Platform for guidance with analysis of the stroke volumes in Imaris and with BrainJ, and for providing the cold light source to induce photothrombotic strokes.

## Author contributions

ACM and RMC conceived of the project and wrote the manuscript. ACM made figures and illustrations. ACM and LJS performed surgeries. ACM, TXC, and LJS performed behavior experiments. ACM and TXC processed and analyzed brain tissue. HFMR and TT developed and built joystick hardware. RH developed TeenScience board and implemented PID control software. ACM and TXC wrote task control code. TXC performed lightning-pose video analysis. ACM, LJS, JNI, and VRA analyzed mouse behavior data. JNI performed the calibration of motor torques and calculated workspace resolution. JMM designed and trained reinforcement learning model and analyzed agent data.

---

**Supplementary Figure 1.**
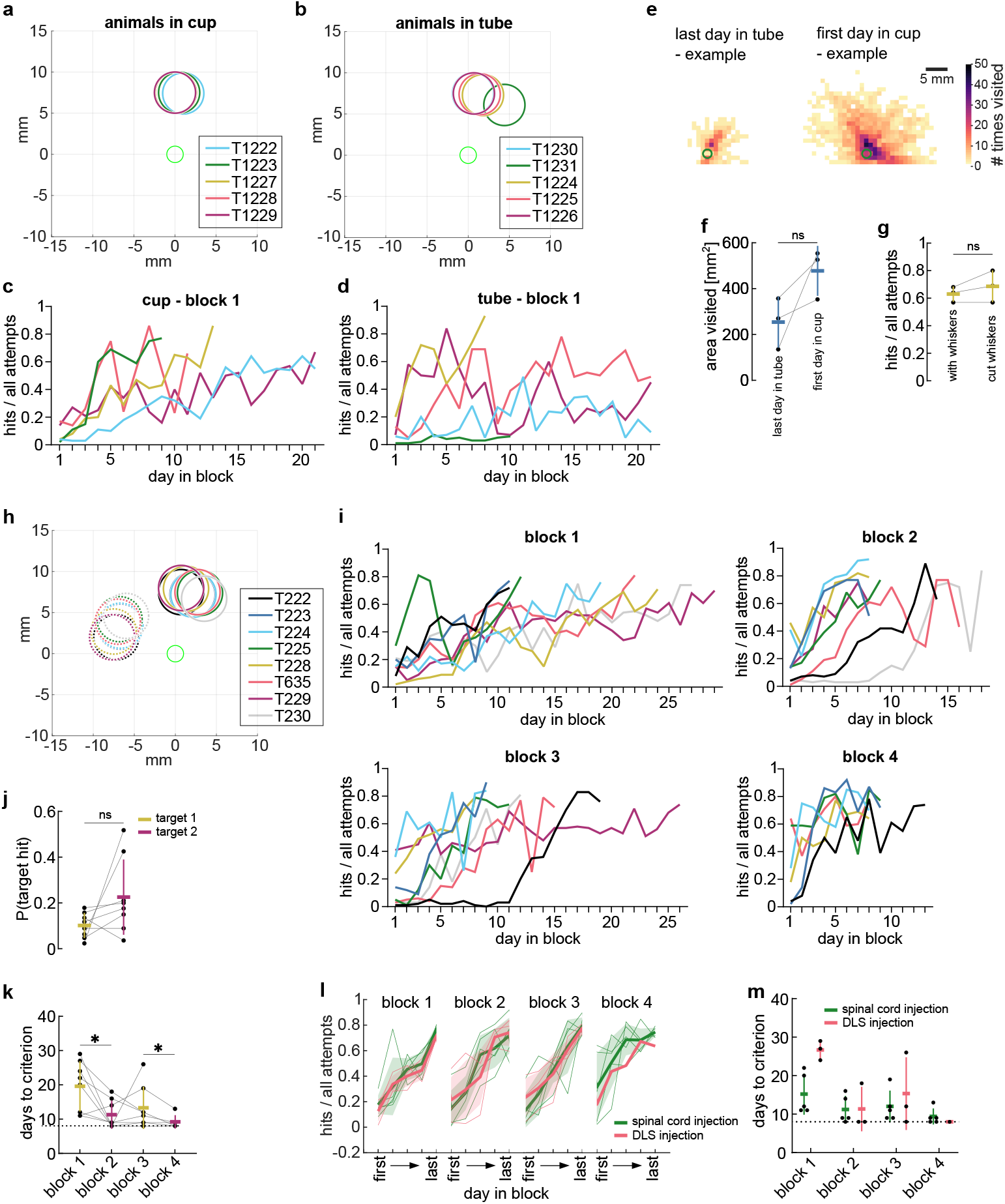
**a-d**. Colors indicate animals trained in the same joystick setup. **a**. Location of targets for animals trained in the cup. **b**. Location of targets for animals trained in the tube. **c**. Hit ratio of animals trained in the cup for all sessions of block 1 until performance or termination criterion was reached. **d**. Same as (c) for animals trained in the tube. **e**. Example heat maps showing the number of times a 1 mm2 bin in the workspace was visited during the last session in the tube and the first session in the cup of the same animal (green circle = start position). **f**. Total area visited by all trajectories on the last day in the tube and the first day in the cup for 3 animals. **g**. Hit ratio of animals that learned to hit target 1 with intact whiskers and on the day after whiskers were cut (paired t-test: t(2) = 1.63, p = 0.245, n = 3). **h**. Location of targets for all animals of the main cohort, target 1 = solid circles, target 2 = dashed circles. **i**. Hit ratio for all sessions until performance criteria for blocks 1-3 (n = 8) and block 4 (n = 6). **j**. Probability of entering defined targets with attempts made on the last day of pre-training. **k**. Number of days in each block to reach the performance criterion showing an overall target effect. **l**. Hit ratio across all blocks of animals grouped by injection into the spinal cord or dorsolateral striatum (DLS) shows no difference in performance (Mixed-effects model, day: F(3.8,20.9) = 12.34, p < 0.0001, group: F(1,6) = 0.72, p = 0.429, spinal cord injection: n = 5, DLS injection: n = 3). **m**. Number of days in each block to reach the performance criterion shows no overall effect of injection location (Mixed-effects model, block: F(3,16) = 7.68, p = 0.002, group: F(1,6) = 2.71, p = 0.151, spinal cord injection: n = 5, DLS injection: n = 3).

**Supplementary Figure 2.**
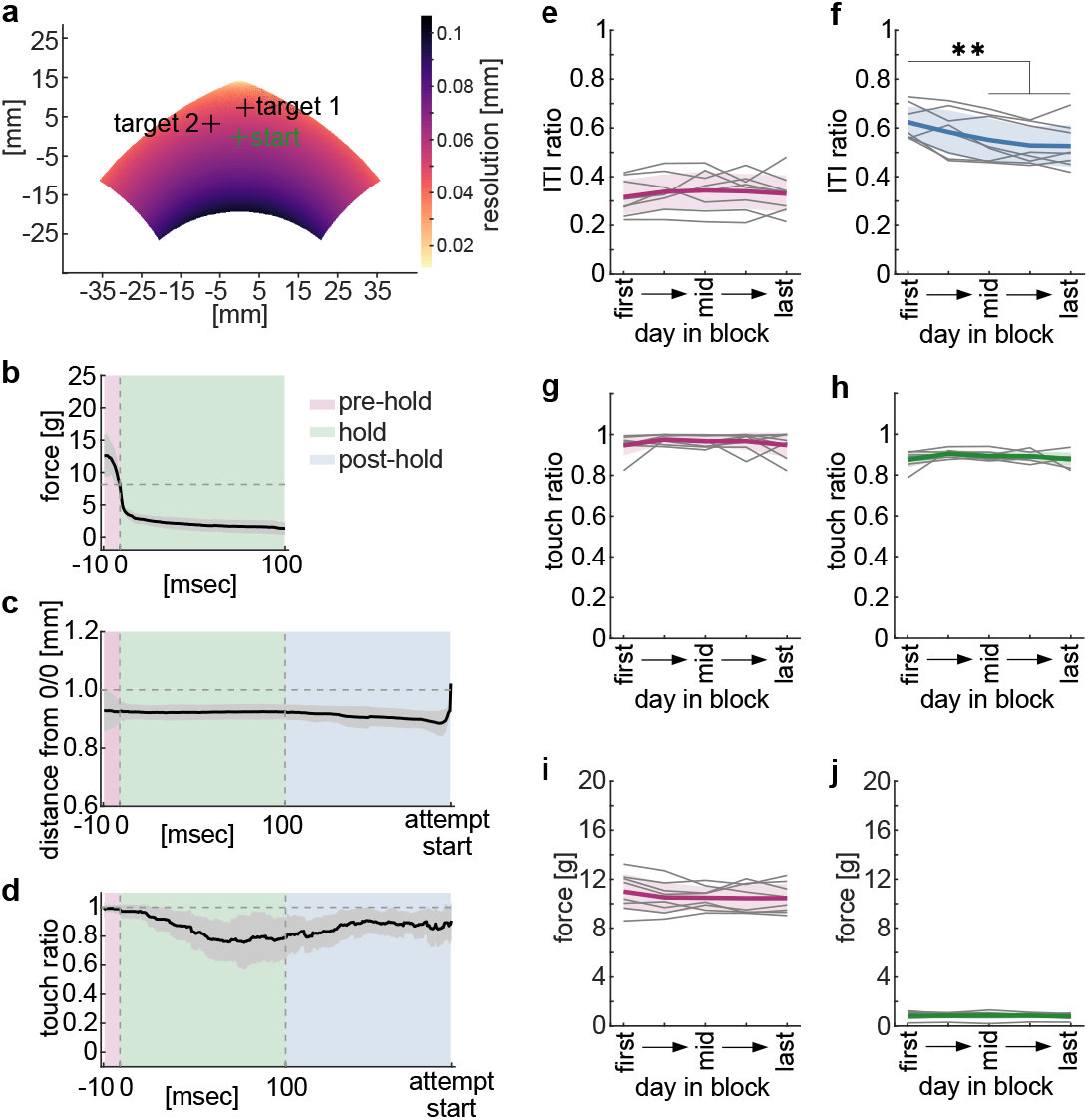
**e-j**. Analysis using all inter-trial-intervals (ITI) on 5 selected days and averaged across all blocks (n = 8 animals). One-way ANOVA with repeated measures, asterisks show Dunnett’s multiple comparisons between the first day and all other days, **: p < 0.0. Mean +/-stdev and single animals. **a**. Workspace and spatial resolution of SCARA joystick and mean target locations. **b**. Force profile across all sessions of an example animal during the pre-hold and hold periods of the ITI. Horizontal dashed line shows the threshold of 7g. The force had to be below this threshold for 100 msec to exit the hold period. **c**. Same example data as in (b) showing the joystick position during the pre-hold, hold and post-hold periods. The post-hold period is resampled for all trials between the end of the hold period and the time point of leaving the start position (end of the ITI). The horizontal dashed line shows the radius of the start position. Crossing the radius initiates an attempt. **d**. Same example data as (b/c) showing the probability of joystick touch during the pre-hold, hold, and post-hold periods. **e**. Ratio of ITIs that included a pre-hold period during which the animal exerted force above the threshold for at least 10 msec (one-way ANOVA, F(2.6,18.0) = 0.73, p = 0.525). **f**. Ratio of ITIs that included a post-hold period during which the animal was not yet leaving the start position after having exited the hold period (one-way ANOVA, F(2.7,18.8) = 13.18, p = 0.0001). **g**. Probability of joystick touch during the pre-hold period (one-way ANOVA, F(2.8,19.5) = 0.82, p = 0.490). **h**. Probability of joystick touch during the hold period (one-way ANOVA, F(1.4,10.1) = 1.89, p = 0.202). **i**. Force exerted by the animal against the joystick during the pre-hold period (one-way ANOVA, F(1.9,13.3) = 1.42, p = 0.275). **j**. Force exerted by the animal against the joystick during the hold period (one-way ANOVA, F(2.1,14.4) = 0.56, p = 0.590).

**Supplementary Figure 3.**
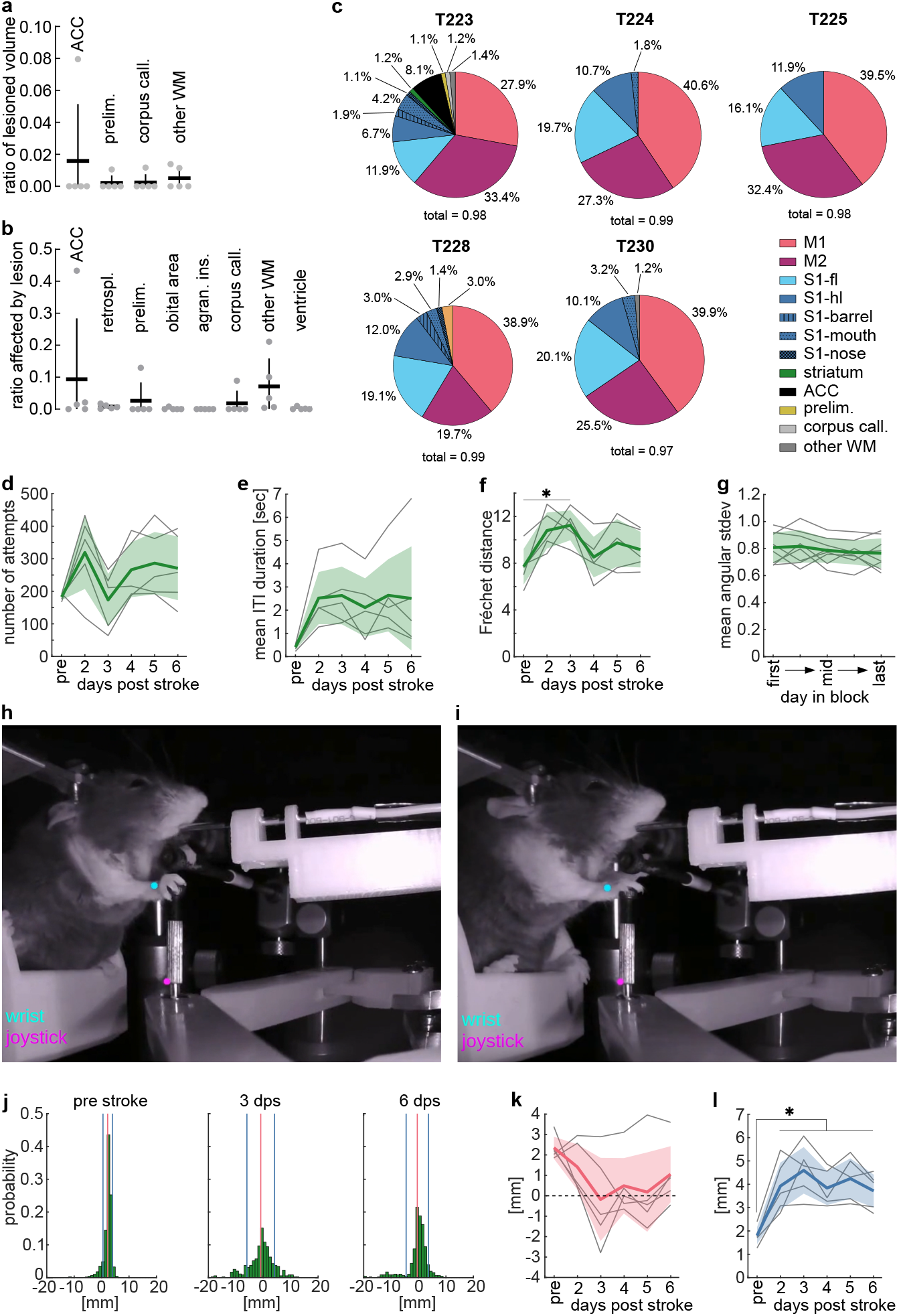
One-way ANOVA with repeated measures, asterisks show Dunnett’s multiple comparisons between the pre-stroke day and all post-stroke days of p < 0.05. Mean +/-stdev and single animals (n = 5). **a**. Relative volume of total stroke lesion that affected different additional Allen Brain Atlas Reference Brain areas. **b**. Additional Allen Brain Atlas Reference Brain areas that were affected by the stroke lesion showing the relative volume lesioned. **c**. Pie charts for individual animals showing relative volume of different Allen Brain Atlas Reference Brain of the total stroke lesion. Only areas comprising > 1% of the total stroke lesion volume are shown **d**. Total number of attempts made per session before and after the stroke lesion. **e**. Mean time between attempts before and after the stroke lesion. **f**. Average similarity between trajectories pre-stroke and post-stroke measured by pair-wise Fréchet distance shows trajectories pre-stroke are more similar to each other than trajectories post-stroke. **g**. Mean spatial variability of full length trajectories of all attempts across all blocks. **h**. Example video frame showing wrist and joystick key points tracked using lightning pose in a pre-stroke session, (cyan = wrist, magenta = joystick). **i**. Same as (h) but 2 days post stroke. **j**. Example data from a single animal pre-stroke and 3 and 6 days post stroke showing histograms of the distance between the wrist and the joystick key points in the x-dimension of the video during active joystick movements. Red line = mean, blue lines = +/-stdev. **k**. Mean x-dimension distance between the wrist and joystick for all animals and sessions. Positive values = the wrist is left of the joystick in the x-dimension of the video frame. Negative values = the wrist is right of the joystick in the x-dimension of the video frame. **l**. Standard deviation of x-dimension distance between the wrist and joystick for all animals and sessions. a-c. Abbreviations. M1: primary motor cortex, M2: secondary motor cortex, S1-fl: primary sensory cortex – forelimb, S1-hl: primary sensory cortex – hindlimb, S1-others: primary sensory cortex – other areas, ACC: anterior cingulate cortex, retrospl.: retrosplenial, prelim.: prelimbic, agran. ins.: agranular insular, corpus call.: corpus callosum, WM: white matter.

**Supplementary Figure 4.**
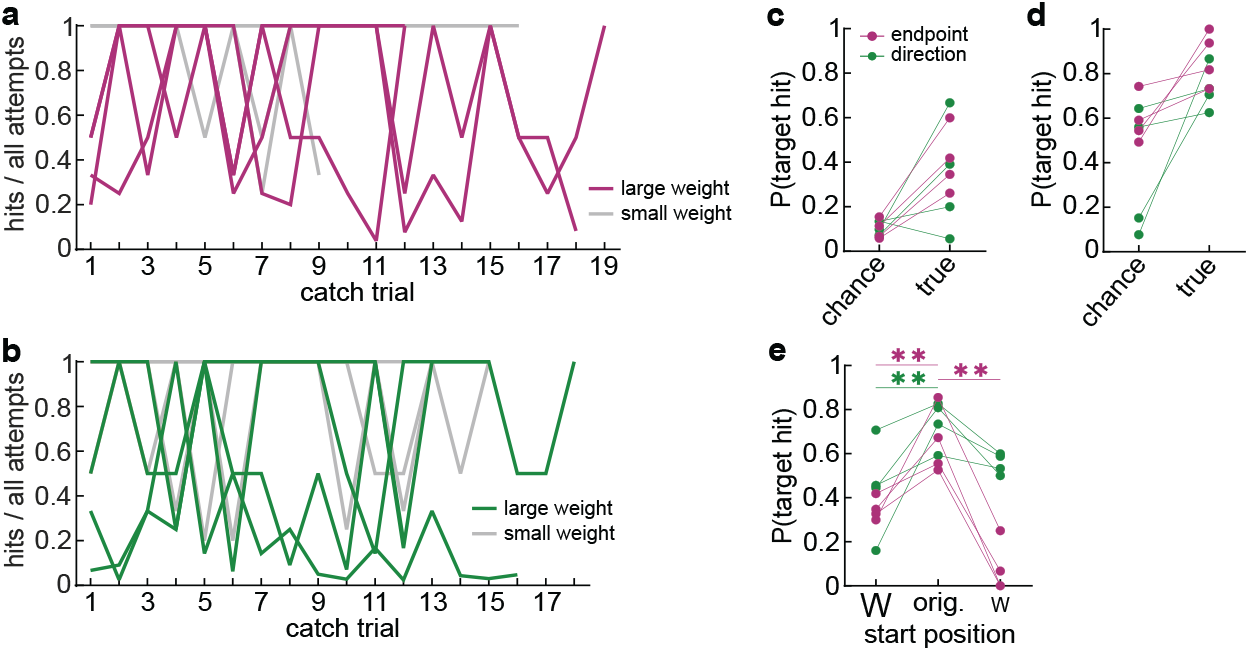
**a**. Hit ratio for each catch trial event for all endpoint learner animals showing the side with the larger weight in color and the side with the smaller weight in grey. **b**. Same as (a) but for direction learner type animals. **c**. Chance and true hit ratio from the start position with a larger weight. Chance hit ratio was calculated by translating all attempts made from the original start to the new start. True hit ratio is the actual mean hit ratio achieved by the animals from the new start position across all catch trials. **d**. Same as (c) but for start with smaller weight. **e**. Probability of target hit for attempts made from new starts translated to original start positions compared to the true hit ratio in from the original start position. Start positions split by size of weight (w).

**Supplementary Table 1.**

| Experiment | Animal ID | Trained in | Number of blocks | Test after block | AAV injection site | AAV | Provider / Lot number | Volume | Segments | Cortex lesion |
| --- | --- | --- | --- | --- | --- | --- | --- | --- | --- | --- |
| Cup and Tube | T1222 | Cup | 1 |  | No injection |  |  |  |  |  |
| Cup and Tube | T1223 | Cup | 1 |  | No injection |  |  |  |  |  |
| Cup and Tube | T1224 | Tube | 1 |  | No injection |  |  |  |  |  |
| Cup and Tube | T1225 | Tube | 1 |  | No injection |  |  |  |  |  |
| Cup and Tube | T1226 | Tube | 1 |  | No injection |  |  |  |  |  |
| Cup and Tube | T1227 | Cup | 1 |  | No injection |  |  |  |  |  |
| Cup and Tube | T1228 | Cup | 1 |  | No injection |  |  |  |  |  |
| Cup and Tube | T1229 | Cup | 1 |  | No injection |  |  |  |  |  |
| Cup and Tube | T1230 | Tube | 1 |  | No injection |  |  |  |  |  |
| Cup and Tube | T1231 | Tube | 1 |  | No injection |  |  |  |  |  |
| Target Training | T222 | Cup | 4 | 5 | Spinal cord | AAVretro(SL1)_Syn_GCamp6f | Janelia | 75 nl/seg. | C4 – C8 | No |
| Target Training | T223 | Cup | 4 | 5 | Spinal cord | AAVretro(SL1)_Syn_GCamp6f | Janelia | 75 nl/seg. | C4 – C8 | Yes |
| Target Training | T224 | Cup | 4 | 5 | Spinal cord | AAVretro(SL1)_Syn_GCamp6f | Janelia | 75 nl/seg. | C4 – C7 | Yes |
| Target Training | T225 | Cup | 4 | 5 | Spinal cord | AAVretro(SL1)_Syn_GCamp6f | Janelia | 75 nl/seg. | C4 – C7 | Yes |
| Target Training | T228 | Cup | 4 | 5 | DLS | AAVretro(SL1)_Syn_GCamp6f | Janelia | 100 nl |  | Yes |
| Target Training | T229 | Cup | 3 | 3 | DLS | AAVretro(SL1)_Syn_GCamp6f | Janelia | 100 nl |  | No |
| Target Training | T230 | Cup | 3 | 3 | DLS | AAVretro(SL1)_Syn_GCamp6f | Janelia | 100 nl |  | Yes |
| Target Training | T635 | Cup | 4 | 3 | Spinal cord | AAVretro_Syn_GCamp7f | 21720 | 75 nl/seg. | C4 – C8 | No |
| Whisker trim | T638 | Cup | 3 | 3 | Spinal cord | AAVretro_Syn_GCamp7f | v52598 | 60 nl/seg. | C4 – C8 |  |
| Whisker trim | T640 | Cup | 3 | 3 | DLS | AAVretro_Syn_GCamp7f | 21720 | 80 nl |  |  |
| Whisker trim | T644 | Cup | 3 | 3 | DLS | AAVretro_Syn_GCamp7f | v52598 | 80 nl |  |  |

**Supplementary Table 2.**

| Figure | Panel | Mixed-effects model day<br>in block effect (F) | Mixed-effects model day<br>in block effect (p-value) | Mixed-effects model block<br>effect (F) | Mixed-effects model block<br>effect (p-value) |
| --- | --- | --- | --- | --- | --- |
| 2 | b | $F(2.6, 18.1) = 14.33$ | 0.0001 | $F(1.7, 12.0) = 3.84$ | 0.057 |
| 2 | c | $F(2.9, 20.4) = 7.13$ | 0.002 | $F(2.4, 16.8) = 0.28$ | 0.797 |
| 2 | f | $F(2.1, 14.8) = 8.46$ | 0.003 | $F(2.1, 14.5) = 0.91$ | 0.428 |
| 2 | h | $F(2.2, 15.3) = 15.31$ | 0.0002 | $F(2.2, 15.0) = 0.72$ | 0.513 |
| 2 | j (top row) | $F(3.1, 21.4) = 4.17$ | 0.018 | $F(1.9, 13.1) = 1.55$ | 0.249 |
| 2 | k / j (diagonal) | $F(2.9, 20.1) = 13.93$ | < 0.0001 | $F(2.1, 14.4) = 0.16$ | 0.862 |
| 2 | m | $F(2.1, 14.4) = 6.17$ | 0.011 | $F(1.8, 12.4) = 0.35$ | 0.689 |
| 2 | n | $F(2.3, 16.0) = 8.67$ | 0.002 | $F(2.2, 15.2) = 0.34$ | 0.735 |
| 2 | o | $F(1.9, 13.4) = 5.22$ | 0.022 | $F(2.3, 15.9) = 1.43$ | 0.269 |
| 2 | p | $F(1.9, 13.0) = 9.60$ | 0.003 | $F(2.4, 16.9) = 0.97$ | 0.414 |
| 3 | b | $F(1.4, 9.7) = 7.54$ | 0.016 | $F(1.6, 11.1) = 2.35$ | 0.147 |
| 3 | c | $F(2.4, 16.8) = 11.43$ | 0.0005 | $F(1.9, 13.1) = 0.22$ | 0.790 |
| 3 | e | $F(2.3, 16.2) = 6.28$ | 0.008 | $F(1.9, 13.1) = 1.65$ | 0.229 |
| 3 | f | $F(1.9, 13.5) = 1.33$ | 0.296 | $F(1.9, 13.5) = 0.30$ | 0.736 |
| 3 | g | $F(3.1, 21.4) = 4.46$ | 0.014 | $F(2.5, 17.5) = 0.48$ | 0.667 |
| 3 | i | $F(2.1, 14.7) = 9.88$ | 0.002 | $F(2.3, 16.4) = 1.76$ | 0.200 |
| 3 | j | $F(2.1, 14.6) = 4.68$ | 0.026 | $F(2.1, 14.8) = 1.82$ | 0.195 |
| Suppl. 2 | e | $F(2.3, 16.0) = 0.60$ | 0.580 | $F(1.5, 10.7) = 1.76$ | 0.219 |
| Suppl. 2 | f | $F(2.7, 18.6) = 9.60$ | 0.0007 | $F(1.4, 9.9) = 3.20$ | 0.095 |
| Suppl. 2 | g | $F(2.5, 17.8) = 0.91$ | 0.442 | $F(1.2, 8.1) = 1.89$ | 0.209 |
| Suppl. 2 | h | $F(1.3, 9.1) = 2.53$ | 0.143 | $F(1.3, 9.1) = 1.07$ | 0.350 |
| Suppl. 2 | i | $F(1.9, 13.6) = 1.14$ | 0.347 | $F(1.7, 11.6) = 2.82$ | 0.107 |
| Suppl. 2 | j | $F(2.2, 15.2) = 0.51$ | 0.623 | $F(1.3, 9.4) = 0.42$ | 0.590 |
| Suppl. 3 | g | $F(2.4, 17.0) = 0.93$ | 0.431 | $F(2.0, 13.7) = 2.08$ | 0.163 |

## REFERENCES

1. A. Raichin, A. Shkedy Rabani, L. Shmuelof, Motor skill training without online visual feedback enhances feedforward control. J Neurophysiol 126, 1604–1613(2021).

2. E. J. Hwang, R. Shadmehr, Internal models of limb dynamics and the encoding of limb state. J Neural Eng 2, S266–278 (2005).

3. M. A. Goodale, D. Pelisson, C. Prablanc, Large adjustments in visually guided reaching do not depend on vision of the hand or perception of target displacement. Nature 320, 748–750 (1986).

4. N. Yousif, J. Diedrichsen, Structural learning in feedforward and feedback control. J Neurophysiol 108, 2373–2382 (2012).

5. S. Kasuga, S. Telgen, J. Ushiba, D. Nozaki, J. Diedrichsen, Learning feedback and feedforward control in a mirror-reversed visual environment. J Neurophysiol 114, 2187–2193 (2015).

6. F. Hoogewoud, A. Hamadjida, A. F. Wyss, A. Mir, M. E. Schwab, A. Belhaj-Saif, E. M. Rouiller, Comparison of functional recovery of manual dexterity after unilateral spinal cord lesion or motor cortex lesion in adult macaque monkeys. Front Neurol 4, 101 (2013).

7. I. Q. Whishaw, Loss of the innate cortical engram for action patterns used in skilled reaching and the development of behavioral compensation following motor cortex lesions in the rat. Neuropharmacology 39, 788–805 (2000).

8. O. A. Gharbawie, C. L. Gonzalez, I. Q. Whishaw, Skilled reaching impairments from the lateral frontal cortex component of middle cerebral artery stroke: a qualitative and quantitative comparison to focal motor cortex lesions in rats. Behav Brain Res 156, 125–137 (2005).

9. W. G. Darling, M. A. Pizzimenti, R. J. Morecraft, Functional recovery following motor cortex lesions in non-human primates: experimental implications for human stroke patients. J Integr Neurosci 10, 353–384 (2011).

10. J. Z. Guo, A. R. Graves, W. W. Guo, J. Zheng, A. Lee, J. Rodriguez-Gonzalez, N. Li, J. J. Macklin, J. W. Phillips, B. D. Mensh, K. Branson, A. W. Hantman, Cortex commands the performance of skilled movement. Elife 4, e10774 (2015).

11. K. Morandell, D. Huber, The role of forelimb motor cortex areas in goal directed action in mice. Sci Rep 7, 15759 (2017).

12. K. A. Tennant, T. A. Jones, Sensorimotor behavioral effects of endothelin-1 induced small cortical infarcts in C57BL/6 mice. J Neurosci Methods 181, 18–26 (2009).

13. R. Caminiti, P. B. Johnson, A. Urbano, Making arm movements within different parts of space: dynamic aspects in the primate motor cortex. J Neurosci 10, 2039–2058 (1990).

14. A. P. Georgopoulos, J. F. Kalaska, R. Caminiti, J. T. Massey, On the relations between the direction of two-dimensional arm movements and cell discharge in primate motor cortex. J Neurosci 2, 1527–1537 (1982).

15. A. P. Georgopoulos, A. B. Schwartz, R. E. Kettner, Neuronal population coding of movement direction. Science 233, 1416–1419 (1986).

16. J. F. Kalaska, From intention to action: motor cortex and the control of reaching movements. Adv Exp Med Biol 629, 139–178 (2009).

17. S. H. Scott, J. F. Kalaska, Changes in motor cortex activity during reaching movements with similar hand paths but different arm postures. J Neurophysiol 73, 2563–2567 (1995).

18. J. Rickert, A. Riehle, A. Aertsen, S. Rotter, M. P. Nawrot, Dynamic encoding of movement direction in motor cortical neurons. J Neurosci 29, 13870–13882 (2009).

19. J. A. Pruszynski, I. Kurtzer, J. Y. Nashed, M. Omrani, B. Brouwer, S. H. Scott, Primary motor cortex underlies multi-joint integration for fast feedback control. Nature 478, 387–390 (2011).

20. S. H. Scott, Optimal feedback control and the neural basis of volitional motor control. Nat Rev Neurosci 5, 532–546 (2004).

21. M. M. Pacheco, C. W. Lafe, K. M. Newell, Search Strategies in the Perceptual-Motor Workspace and the Acquisition of Coordination, Control, and Skill. Front Psychol 10, 1874 (2019).

22. P. Redgrave, K. Gurney, The short-latency dopamine signal: a role in discovering novel actions? Nat Rev Neurosci 7, 967–975 (2006).

23. M. S. Fee, The role of efference copy in striatal learning. Curr Opin Neurobiol 25, 194–200 (2014).

24. L. Shmuelof, J. W. Krakauer, P. Mazzoni, How is a motor skill learned? Change and invariance at the levels of task success and trajectory control. J Neurophysiol 108, 578–594 (2012).

25. E. Todorov, M. I. Jordan, Optimal feedback control as a theory of motor coordination. Nat Neurosci 5, 1226–1235 (2002).

26. E. Thorndike, Some Experiments on Animal Intelligence. Science 7, 818–824 (1898).

27. F. J. Santos, R. F. Oliveira, X. Jin, R. M. Costa, Corticostriatal dynamics encode the refinement of specific behavioral variability during skill learning. Elife 4, e09423 (2015).

28. T. Stafford, M. Thirkettle, T. Walton, N. Vautrelle, L. Hetherington, M. Port, K. Gurney, P. Redgrave, A novel task for the investigation of action acquisition. PLoS One 7, e37749 (2012).

29. A. P. Georgopoulos, J. F. Kalaska, J. T. Massey, Spatial trajectories and reaction times of aimed movements: effects of practice, uncertainty, and change in target location. J Neurophysiol 46, 725–743 (1981).

30. A. K. Dhawale, Y. R. Miyamoto, M. A. Smith, B. P. Olveczky, Adaptive Regulation of Motor Variability. Curr Biol 29, 3551–3562 e3557 (2019).

31. S. E. Pekny, J. Izawa, R. Shadmehr, Reward-dependent modulation of movement variability. J Neurosci 35, 4015–4024 (2015).

32. H. G. Wu, Y. R. Miyamoto, L. N. Gonzalez Castro, B. P. Olveczky, M. A. Smith, Temporal structure of motor variability is dynamically regulated and predicts motor learning ability. Nat Neurosci 17, 312–321 (2014).

33. C. L. Hull, Principles of behavior: an introduction to behavior theory. (Appleton-Century, 1943).

34. E. C. Tolman, B. F. Ritchie, D. Kalish, Studies in spatial learning: Orientation and the short-cut. J Exp Psychol 36, 13–24 (1946).

35. E. C. Tolman, B. F. Ritchie, D. Kalish, Studies in spatial learning; place learning versus response learning. J Exp Psychol 36, 221–229 (1946).

36. R. Morris, Developments of a water-maze procedure for studying spatial learning in the rat. J Neurosci Methods 11, 47–60 (1984).

37. C. D. Adams, A. Dickinson, Instrumental Responding Following Reinforcer Devaluation. Q J Exp Psychol-B 33, 109–121 (1981).

38. J. W. Krakauer, Z. M. Pine, M. F. Ghilardi, C. Ghez, Learning of visuomotor transformations for vectorial planning of reaching trajectories. J Neurosci 20, 8916–8924 (2000).

39. N. Malfait, D. M. Shiller, D. J. Ostry, Transfer of motor learning across arm configurations. J Neurosci 22, 9656–9660 (2002).

40. R. Shadmehr, Z. M. Moussavi, Spatial generalization from learning dynamics of reaching movements. J Neurosci 20, 7807–7815 (2000).

41. R. Shadmehr, F. A. Mussa-Ivaldi, Adaptive representation of dynamics during learning of a motor task. J Neurosci 14, 3208–3224 (1994).

42. J. B. Brayanov, D. Z. Press, M. A. Smith, Motor memory is encoded as a gain-field combination of intrinsic and extrinsic action representations. J Neurosci 32, 14951–14965 (2012).

43. A. Polit, E. Bizzi, Characteristics of motor programs underlying arm movements in monkeys. J Neurophysiol 42, 183–194 (1979).

44. E. Bizzi, N. Accornero, W. Chapple, N. Hogan, Posture control and trajectory formation during arm movement. J Neurosci 4, 2738–2744 (1984).

45. M. Thirkettle, T. Walton, A. Shah, K. Gurney, P. Redgrave, T. Stafford, The path to learning: action acquisition is impaired when visual reinforcement signals must first access cortex. Behav Brain Res 243, 267–272 (2013).

46. O. Lambercy, M. Schubring-Giese, B. Vigaru, R. Gassert, A. R. Luft, J. A. Hosp, Sub-processes of motor learning revealed by a robotic manipulandum for rodents. Behav Brain Res 278, 569–576 (2015).

47. B. Vigaru, O. Lambercy, L. Graber, R. Fluit, P. Wespe, M. Schubring-Giese, A. Luft, R. Gassert, A small-scale robotic manipulandum for motor training in stroke rats. IEEE Int Conf Rehabil Robot 2011, 5975349 (2011).

48. B. C. Vigaru, O. Lambercy, M. Schubring-Giese, J. A. Hosp, M. Schneider, C. Osei-Atiemo, A. Luft, R. Gassert, A robotic platform to assess, guide and perturb rat forelimb movements. IEEE Trans Neural Syst Rehabil Eng 21, 796–805 (2013).

49. M. J. Wagner, J. Savall, T. H. Kim, M. J. Schnitzer, L. Luo, Skilled reaching tasks for head-fixed mice using a robotic manipulandum. Nat Protoc 15, 1237–1254 (2020).

50. T. Bollu, S. C. Whitehead, N. Prasad, J. Walker, N. Shyamkumar, R. Subramaniam, B. Kardon, I. Cohen, J. H. Goldberg, Automated home cage training of mice in a hold-still center-out reach task. J Neurophysiol 121, 500–512 (2019).

51. B. Panigrahi, K. A. Martin, Y. Li, A. R. Graves, A. Vollmer, L. Olson, B. D. Mensh, A. Y. Karpova, J. T. Dudman, Dopamine Is Required for the Neural Representation and Control of Movement Vigor. Cell 162, 1418–1430 (2015).

52. J. Park, J. W. Phillips, J. Z. Guo, K. A. Martin, A. W. Hantman, J. T. Dudman, Motor cortical output for skilled forelimb movement is selectively distributed across projection neuron classes. Sci Adv 8, eabj5167 (2022).

53. M. J. Wagner, J. Savall, O. Hernandez, G. Mel, H. Inan, O. Rumyantsev, J. Lecoq, T. H. Kim, J. Z. Li, C. Ramakrishnan, K. Deisseroth, L. Luo, S. Ganguli, M. J. Schnitzer, A neural circuit state change underlying skilled movements. Cell 184, 3731–3747 e3721 (2021).

54. M. J. Wagner, T. H. Kim, J. Kadmon, N. D. Nguyen, S. Ganguli, M. J. Schnitzer, L. Luo, Shared Cortex-Cerebellum Dynamics in the Execution and Learning of a Motor Task. Cell 177, 669–682 e624 (2019).

55. M. J. Wagner, T. H. Kim, J. Savall, M. J. Schnitzer, L. Luo, Cerebellar granule cells encode the expectation of reward. Nature 544, 96–100 (2017).

56. K. A. Tennant, A. L. Asay, R. P. Allred, A. R. Ozburn, J. Kleim, T. A. Jones, The vermicelli and capellini handling tests: simple quantitative measures of dexterous forepaw function in rats and mice. J Vis Exp, (2010).

57. E. J. Hwang, J. E. Dahlen, Y. Y. Hu, K. Aguilar, B. Yu, M. Mukundan, A. Mitani, T. Komiyama, Disengagement of motor cortex from movement control during long-term learning. Sci Adv 5, eaay0001 (2019).

58. Z. V. Guo, S. A. Hires, N. Li, D. H. O’Connor, T. Komiyama, E. Ophir, D. Huber, C. Bonardi, K. Morandell, D. Gutnisky, S. Peron, N. L. Xu, J. Cox, K. Svoboda, Procedures for behavioral experiments in head-fixed mice. PLoS One 9, e88678 (2014).

59. M. W. Mathis, A. Mathis, N. Uchida, Somatosensory Cortex Plays an Essential Role in Forelimb Motor Adaptation in Mice. Neuron 93, 1493–1503 e1496 (2017).

60. A. J. Peters, S. X. Chen, T. Komiyama, Emergence of reproducible spatiotemporal activity during motor learning. Nature 510, 263–267 (2014).

61. Z. Danziger. (MATLAB Central File Exchange, 2023), vol. 2023.

62. M. Schubring-Giese, K. Molina-Luna, B. Hertler, M. M. Buitrago, D. F. Hanley, A. R. Luft, Speed of motor re-learning after experimental stroke depends on prior skill. Exp Brain Res 181, 359–365 (2007).

63. R. Kawai, T. Markman, R. Poddar, R. Ko, A. L. Fantana, K. Dhawale, A. R. Kampff, B. P. Olveczky, Motor cortex is required for learning but not for executing a motor skill. Neuron 86, 800–812 (2015).

64. V. Labat-gest, S. Tomasi, Photothrombotic ischemia: a minimally invasive and reproducible photochemical cortical lesion model for mouse stroke studies. J Vis Exp, (2013).

65. P. Botta, A. Fushiki, A. M. Vicente, L. A. Hammond, A. C. Mosberger, C. R. Gerfen, D. Peterka, R. M. Costa, An Amygdala Circuit Mediates Experience-Dependent Momentary Arrests during Exploration. Cell 183, 605–619 e622 (2020).

66. A. K. Dhawale, M. A. Smith, B. P. Olveczky, The Role of Variability in Motor Learning. Annu Rev Neurosci 40, 479–498 (2017).

67. M. M. Pacheco, K. M. Newell, Transfer as a function of exploration and stabilization in original practice. Hum Mov Sci 44, 258–269 (2015).

68. W. D. Beggs, C. I. Howarth, The movement of the hand towards a target. Q J Exp Psychol 24, 448–453 (1972).

69. T. A. Martin, J. G. Keating, H. P. Goodkin, A. J. Bastian, W. T. Thach, Throwing while looking through prisms. I. Focal olivocerebellar lesions impair adaptation. Brain 119 (Pt 4), 1183–1198 (1996).

70. D. M. Wolpert, R. C. Miall, M. Kawato, Internal models in the cerebellum. Trends Cogn Sci 2, 338–347 (1998).

71. D. M. Wolpert, Z. Ghahramani, M. I. Jordan, An internal model for sensorimotor integration. Science 269, 1880–1882 (1995).

72. R. Shadmehr, M. A. Smith, J. W. Krakauer, Error correction, sensory prediction, and adaptation in motor control. Annu Rev Neurosci 33, 89–108 (2010).

73. D. J. Palidis, J. G. A. Cashaback, P. L. Gribble, Neural signatures of reward and sensory error feedback processing in motor learning. J Neurophysiol 121, 1561–1574 (2019).

74. J. Izawa, R. Shadmehr, Learning from sensory and reward prediction errors during motor adaptation. PLoS Comput Biol 7, e1002012 (2011).

75. A. T. Welford, A. H. Norris, N. W. Shock, Speed and accuracy of movement and their changes with age. Acta Psychol (Amst) 30, 3–15 (1969).

76. D. Sternad, It’s Not (Only) the Mean that Matters: Variability, Noise and Exploration in Skill Learning. Curr Opin Behav Sci 20, 183–195 (2018).

77. A. Renart, C. K. Machens, Variability in neural activity and behavior. Curr Opin Neurobiol 25, 211–220 (2014).

78. A. A. Faisal, L. P. Selen, D. M. Wolpert, Noise in the nervous system. Nat Rev Neurosci 9, 292–303 (2008).

79. C. M. Harris, D. M. Wolpert, Signal-dependent noise determines motor planning. Nature 394, 780–784 (1998).

80. A. S. Therrien, D. M. Wolpert, A. J. Bastian, Increasing Motor Noise Impairs Reinforcement Learning in Healthy Individuals. eNeuro 5, p(2018).

81. E. C. Tumer M. S. Brainard, Performance variability enables adaptive plasticity of ‘crystallized’ adult birdsong. Nature 450, 1240–1244 (2007).

82. M. H. Kao, M. S. Brainard, Lesions of an avian basal ganglia circuit prevent context-dependent changes to song variability. J Neurophysiol 96, 1441–1455 (2006).

83. S. T. Albert, R. Shadmehr, The Neural Feedback Response to Error As a Teaching Signal for the Motor Learning System. J Neurosci 36, 4832–4845 (2016).

84. I. S. Alonso, I.; Palacio-Manzano, M.; Frézel-Jacob, N.; Philippides A., Peripersonal encoding of forelimb proprioception in the mouse somatosensory cortex. BioRxiv, (2023).

85. G. L. Galinanes, C. Bonardi, D. Huber, Directional Reaching for Water as a Cortex-Dependent Behavioral Framework for Mice. Cell Rep 22, 2767–2783 (2018).

86. A. J. Peters, H. Liu, T. Komiyama, Learning in the Rodent Motor Cortex. Annu Rev Neurosci 40, 77–97 (2017).

87. J. A. Kleim, S. Barbay, R. J. Nudo, Functional reorganization of the rat motor cortex following motor skill learning. J Neurophysiol 80, 3321–3325 (1998).

88. A. R. Luft, M. M. Buitrago, T. Ringer, J. Dichgans, J. B. Schulz, Motor skill learning depends on protein synthesis in motor cortex after training. J Neurosci 24, 6515–6520 (2004).

89. B. A. Sauerbrei, J. Z. Guo, J. D. Cohen, M. Mischiati, W. Guo, M. Kabra, N. Verma, B. Mensh, K. Branson, A. W. Hantman, Cortical pattern generation during dexterous movement is input-driven. Nature 577, 386–391 (2020).

90. S. B. E. Wolff, R. Ko, B. P. Olveczky, Distinct roles for motor cortical and thalamic inputs to striatum during motor skill learning and execution. Sci Adv 8, eabk0231 (2022).

91. M. D. Golub, B. M. Yu, A. B. Schwartz, S. M. Chase, Motor cortical control of movement speed with implications for brain-machine interface control. J Neurophysiol 112, 411–429 (2014).

92. D. W. Biderman, M.R.; Hurwitz, C.; Greenspan, N.R.; Lee, R.S.; Vishnubhotla, A.; Schartner, M.; Huntenburg, J.M.; Khanal, A.; Meijer, G.T.; Noel, J-P.; Pan-Vazquez, A.; Socha K.Z.; Urai, A.E.; The International Brain Laboratory, Warren, R.; Noone, D.; Pedraja, F.; Cunningham, J.; Sawtell, N.B.; Paninski, L., Lightning Pose: improved animal pose estimation via semi-supervised learning, Bayesian ensembling, and cloud-native opensource tools. BioRXiv, (2023).

93. T. Abe, I. Kinsella, S. Saxena, E. K. Buchanan, J. Couto, J. Briggs, S. L. Kitt, R. Glassman, J. Zhou, L. Paninski, J. P. Cunningham, Neuroscience Cloud Analysis As a Service: An open-source platform for scalable, reproducible data analysis. Neuron 110, 2771–2789 e2777 (2022).

94. T. Walton, M. Thirkettle, P. Redgrave, K. N. Gurney, T. Stafford, The discovery of novel actions is affected by very brief reinforcement delays and reinforcement modality. J Mot Behav 45, 351–360 (2013).

95. D. R. Yatsenko R.; Ecker, A.S.; Walker, E.Y.; Sinz, F.; Berens, P.; Hoenselaar, A.; Cotton, R.J.; Siapas, A.S.; Tolias, A.S., DataJoint: managing big scientific data using MATLAB or Python. BioRXiv, (2015).

96. P. Berens, CircStat: A MATLAB Toolbox for Circular Statistics. Journal of Statistical Software 31, 1–21 (2009)

